# Mapping the living mouse brain neural architecture: strain specific patterns of brain structural and functional connectivity

**DOI:** 10.1101/730366

**Authors:** Meltem Karatas, Vincent Noblet, Md Taufiq Nasseef, Thomas Bienert, Marco Reisert, Jürgen Hennig, Ipek Yalcin, Brigitte Lina Kieffer, Dominik von Elverfeldt, Laura-Adela Harsan

## Abstract

Mapping the structural and functional brain connectivity fingerprints became an essential approach in neurology and experimental neuroscience because network properties can underlie behavioral phenotypes. In mouse models, revealing strain related patterns of brain wiring have a tremendous importance, since these animals are used to answer questions related to neurological or neuropsychiatric disorders. C57BL/6 and BALB/cJ inbred strains are primary “genetic backgrounds” for brain disease modelling and for testing therapeutic approaches. Nevertheless, extensive literature describes basal differences in the behavioral, neuroanatomical and neurochemical profiles of the two strains, which raises the question whether the observed effects are pathology specific or depend on the genetic background. Here we performed a systematic comparative exploration of brain structure and function of C57BL/6 and BALB/cJ mice via Magnetic Resonance Imaging (MRI). We combined voxel-based morphometry (VBM), diffusion MRI and high resolution fiber mapping (hrFM) and resting state functional MRI (rs-fMRI) and depicted brain-wide dissimilarities in the morphology and “connectome” features in the two strains. Particularly C57BL/6 animals show bigger and denser frontal cortical areas, cortico-striatal tracts and thalamic and midbrain pathways, and higher density of fibers in the genu and splenium of the corpus callosum. These features are fairly reflected in the functional connectograms that emphasize differences in “hubness”, frontal cortical and basal forbrain connectivity. We demonstrate strongly divergent reward-aversion circuitry patterns and some variations of the default mode network features. Inter-hemispherical functional connectivity showed flexibility and adjustment regarding the structural patterns in a strain specific manner. We further provide high-resolution tractograms illustrating also inter-individual variability across inter-hemispherical callosal pathways in the BALB/cJ strain.

## 1 Introduction

Animal models of disease - essential for neuroscience research, are gaining huge importance in the recent years with the development of genetically modified mice and novel methods to investigate brain function. However, the diversity in the genetic background of the organisms used for modelling pathology raises questions of the impact of genetic confound on findings (Crawley et al., 1997; Keane et al., 2011). Indeed, each strain of mice display distinct sets of behavioral, neurochemical and neuroanatomical features (Grubb et al., 2004). Among these strains, C57BL/6 and BALB/cJ the commonly used inbred mouse strains, are prime examples of such divergent phenotypes (Anderzhanova et al., 2013; Belzung and Griebel, 2001; Calcagno et al., 2007).

At behavioral level, BALB/cJ strain displays neophobia (i.e. reluctance to engage with novel stimuli) and elevated anxiety (Anderzhanova et al., 2013; Ohl et al., 2001), in contrast to C57BL/6 mice. At young ages, social proximity is not rewarding for BALB/cJ mice (Panksepp and Lahvis, 2007) and they typically show reduced sociability towards conspecifics (Brodkin, 2007; Brodkin et al., 2004; Jacome et al., 2011; Moy et al., 2007; Sankoorikal et al., 2006). Such differences in social behavior are accompanied by differences in early maternal care (Lassi and Tucci, 2017) as BALB/cJ mice present a weaker maternal attachment to offspring compared to C57BL/6 mice. To divulge the underlying mechanisms of different behavioral phenotypes in C57BL/6 and BALB/cJ mice, neurochemical and neuroanatomical studies are indeed necessary (Belzung and Griebel, 2001). For instance, it has been previously demonstrated that serotonergic and dopaminergic neurotransmitter systems show strain-dependent variations in the brain (Calcagno et al., 2007; Guzzetti et al., 2008). At anatomical and microstructural level, longitudinal in-vivo diffusion tensor imaging (DTI) (Le Bihan, 2014; Le Bihan and Breton, 1985) of BALB/cJ and C57BL/6J mouse brains (Kumar et al., 2012) showed as well differences in developmental trajectories along major white matter (WM) tracts, such as the corpus callosum, as well as of gray matter (GM) regions including thalamic areas and frontal motor cortices.

Besides inter-strain differences, within strain inter-individual variations were also reported, especially concerning the BALB/cJ animals. Neuroanatomical studies using histological methods revealed variability of the size - even complete absence - of the corpus callosum (cc) in BALB/cJ mice (Wahlsten, 1974). The size of this structure (Fairless et al., 2008) and whole brain size (Fairless et al., 2012) were correlated with social measures in 30-day-old BALB/cJ mice. In addition to histology, anatomical magnetic resonance imaging (MRI) and DTI of the brain - evaluating morphology and microstructural features (Kim et al., 2012; Kumar et al., 2012)– further revealed associations between sociability and DTI derived parameters in several brain regions (Kim et al., 2012) - including gray matter - of BALB/cJ male mice. For instance, positive regression between fractional anisotropy (FA), an index of water diffusion orientation and indirectly of tissue organization, and social sniffing times was found in regions located in thalamic nuclei, zona incerta, substantia nigra, visual/orbital/ somatosensory cortices and entorhinal cortex of BALB/cJ strain (Kumar et al., 2012).

In clinics, alterations in the corpus callosum region has also been detected in subgroups of autism spectrum disorders (ASD) patients (Alexander et al., 2007; Just et al., 2006). Indeed, behavioral phenotype of BALB/cJ mice also share similarities with certain aspects of ASD, namely the low sociability, higher anxiety and aggression (Brodkin, 2007; Liska and Gozzi, 2016). Such behavioral phenotypes might stem from the underlying architecture of neural circuitries and established patterns of brain communication. Therefore, mapping the brain connectivity blueprints is an essential field of study in neurology and experimental neuroscience.

A detailed non-invasive insight into the whole brain axonal connectivity in vivo has become possible with the development of diffusion based tractography (Horsfield and Jones, 2002) and high-resolution fiber mapping (hrFM) (Calamante et al., 2011; Harsan et al., 2013). In mice, non-invasive DTI based fiber mapping revealed the spatial organization of axonal and dendritic networks in brain WM regions, in good agreement with the spatial projection patterns visualized using viral axonal tracer data (Wu and Zhang, 2016).

Meanwhile, the functional brain communication or connectivity (FC) can be deciphered non-invasively via resting-state functional MRI (rs-fMRI) (Biswal et al., 1995; Chuang and Nasrallah, 2017).

In mouse models, deciphering strain-related patterns in the brain structural and functional connectivity or intra-strain variations of the brain wiring schemes have a tremendous importance, since these animals are used to answer questions relating to human neurological or neuropsychiatric disorders. In this context, the principal goal of our study was to bridge this gap by systematically probing the brain connectivity patterns of two mouse strains extensively used in preclinical neuroscience, including brain connectivity studies: C57BL/6N and BALB/cJ (Grandjean et al., 2017; Grubb et al., 2004; Hübner et al., 2017; Liska et al., 2015; Mechling et al., 2016; Shah et al., 2016a). We thus performed comparative MRI exploration of brain structure and function via a combination of brain voxel-based morphometry (VBM), DTI and high resolution fiber mapping (hrFM) and resting state functional MRI (rs-fMRI) in female animals.

## 2 Materials and Methods

### 2.1 Ethics statement

All procedures were conducted in compliance with the European Directive 2010/63/EU on the protection of animals used for scientific purposes, national guidelines of the German animal protective law for the use of laboratory animals and with permission of the responsible local authorities for the University Medical Center Freiburg (Regierungspräsidium Freiburg, permit numbers: G-08/15).

### 2.2 Animal setup

8-9 weeks old female C57BL/6N (n=11) and BALB/cJ (n=14) mice, purchased from Charles River Laboratories (Sulzfeld, Germany) were used for the MRI experiments. Animals were housed under standard animal room conditions (temperature 21°C, humidity 55-60%, food and water were given ad libitum).

For the rs-fMRI scans, the animals were initially anesthetized moderately using a subcutaneous (s.c.) medetomidine (MD, Domitor, Pfizer, Karlsruhe, Germany) bolus injection of 0.3 mg per kg body weight in 100 μl 0.9% sodium chloride solution. Preparation for the imaging session included the placement of the animal into the adapted imaging bed, stereotactic fixation of the head and attachment of physiological monitoring devices (pulse oximeter clipped to the hind paw, rectal temperature probe and pressure sensitive respiration pad placed underneath the abdomen). 15 minutes after the MD bolus injection, a continuous s.c. MD infusion of 0.6 mg per kg body weight in 200 μl per hour was applied to the animals throughout the rs-fMRI exams. Animals were monitored strictly for optimal physiological conditions (spO2: 97-99%, body temperature: 35.5±1.5°C, respiratory rates: 100-130 breaths/min). For the morphological T2-weighted and diffusion MRI acquisitions, anesthesia was switched to 1.5% isoflurane (Forene; Abbvie Deutschland GmbH & Co. KG, Wiesbaden, Germany) in 1.2 l/min oxygen and respiratory gating was performed. At the end of experiments the mice spontaneously recovered.

### 2.3 Data acquisition

All scans were performed using a 7T small bore animal scanner (Biospec 70/20, Bruker, Ettlingen, Germany), a cryogenically cooled quadrature mouse brain resonator (MRI CryoProbe, Bruker, Ettlingen, Germany) and the ParaVision software version 5.1 (Bruker, Ettlingen, Germany).

Resting-state fMRI data was acquired with T2*-weighted single shot Gradient Echo-echo planar imaging (GE-EPI) sequence using an echo time (TE) of 10 ms and repetition time (TR) of 1700 ms. 12 axial slices of 0.7 mm thickness were positioned on the mouse brain excluding olfactory bulb and cerebellum, field of view (FOV) and acquisition matrix were 19 × 12 mm^2^ and 128 × 80, respectively. A resolution of 0.15 × 0.15 × 0.7 mm^3^ was achieved and 200 volumes were recorded in an interlaced fashion for each run.

High resolution T2-weighted anatomical images (resolution of 0.051 × 0.051 × 0.3 mm^3^) were acquired with a RARE sequence using the following parameters: TE/TR = 50 ms/6514 ms; 48 slices, 0.3 mm slice thickness, interlaced sampling, RARE factor of 4, 2 averages; an acquisition matrix of 256 × 196 and FOV of 1.3×1 cm^2^.

Structural connectivity was investigated based on DTI data. Acquisitions were carried-out using a 4-shot DT-EPI sequence (TE/TR = 26 ms / 7750 ms; gradient duration (δ) = 4 ms and gradient separation (Δ) = 14 ms), with diffusion gradients applied along 30 nonlinear directions (Jones, 2004) and a b-factor of 1000 s/mm^2^. 25 axial slices with 0.5 mm thickness were acquired with a FOV of 1.5 x 1.2 cm² and an acquisition matrix of 160 x 128 resulting in an image resolution of 0.094 x 0.094 x 0.5 mm^3^.

### 2.4 Data processing

#### 2.4.1 Anatomical MRI analysis

T2-weighted images were compared across groups using a VBM framework. First, each image was corrected for bias field inhomogeneity using N4ITK (Tustison et al., 2010). Then, all images were jointly registered in a deformable way using the group-wise registration procedure implemented in the ANTs registration toolbox (http://stnava.github.io/ANTs/) (Avants et al., 2011). Jacobian maps were computed for each estimated deformation field and a logarithmic transformation was applied on them so that dilation (i.e. jacobian greater than one) and contraction (i.e. jacobian between 0 and 1) are mapped on [0; +∞ [and]-∞; 0], respectively. These maps were finally smoothed using a Gaussian kernel (FWHM: 0.2 mm). Intergroup comparisons were conducted at the voxel level using the general linear model implemented in SPM12 (http://www.fil.ion.ucl.ac.uk/spm/). Statistical maps were thresholded at P< 0.05 after false discovery rate (FDR) p-value correction. Results were superimposed on the average morphological image built from the group-wise registration procedure.

#### 2.4.2 Diffusion Tensor MRI data analysis and high-resolution fiber mapping (hrFM)

Post-processing of the diffusion data was performed using an in-house developed DTI and FiberTool software package for SPM (Harsan et al., 2013; Reisert et al., 2011). Diffusion based parameter maps were generated, including FA, mean diffusivity (MD), radial (RD) and axial diffusivities (AD). We further performed diffusion tractography using a global fiber tracking algorithm (Harsan et al., 2013; Reisert et al., 2011). With this family of tractography algorithms, the reconstructed fibers are built with small line segments (particle) described by a spatial position and orientation. These segments are the basic building blocks of a fiber model, bounding together to form the individual fibers. Their orientation and number are adjusted simultaneously and the connections between segments are formed based on a probabilistic procedure (Reisert et al., 2011). To generate fiber density (FD) maps, the number of tracts in each element of a grid was calculated from whole-mouse brain fibers in a manner very similar to previously published methodology (Calamante et al., 2011; Harsan et al., 2013). Further, the method used the continuity information contained in the fibers reconstructed during the global tracking procedure, to introduce sub-voxel information based on supporting information from neighboring voxels. After the generation of sufficient number of fibers passing a voxel at different spatial locations, their density was used as intravoxel information to construct highly resolved spatial histograms of diffusion orientations referred to as hrFM (Video S1). The directionality of the fibers was therefore incorporated into the hrFM by assigning red/green/blue color to different spatial directions: red: mediolateral, green: dorsoventral and blue: rostro-caudal orientation. The grid size was tailored to generate maps of 12 x 12 x 50 µm^3^ of resolution, 8 times the resolution of the acquired data. hrFM allowed individual assessment of the connectivity features, highlighting fine details of the structural brain scaffolding (Video S1).

Diffusion MRI parametric maps were also compared across groups using a voxel-based analysis. The FA and FD maps were jointly registered using the multimodal group-wise registration procedure implemented in the ANTs registration toolbox (Avants et al., 2011). The analysis was then conducted using the voxel-based quantification (VBQ) method (Draganski et al., 2011) which implements a combined weighting/smoothing procedure, avoiding parameter value changes by Gaussian smoothing applied in standardized space. Furthermore, we conducted voxel-wise analysis on FD maps modulated by Jacobian values - in a similar way to the modulated grey matter probability map used in VBM (Good et al., 2001)-in order to quantify the amount of fibers in the standardized space which accurately reflects that of the native space. A Gaussian kernel with a FWHM of 0.5 mm was applied here. Intergroup comparisons were conducted at the voxel level using the general linear model in SPM12. Statistical maps were thresholded at P< 0.05 after false discovery rate (FDR) p-value correction. Results showing higher statistical values for each group were superimposed on the average FD or FA images built from the corresponding group images.

#### 2.4.3 Resting-state fMRI analysis

Each of the resting-state fMRI image volumes were first realigned on the corresponding first scan, using a least square approach and a 6-parameter rigid body transformation in space. The rs-fMRI data was next registered to a morphological template created from individual T2*-weighted scans with ANTs software using deformable SyN algorithm (Avants et al., 2011). Spatially normalized images were smoothed using a Gaussian kernel with FWHM of 0.3×0.3×0.7 mm^3^ on SPM8 and a zero-phase band-pass filter was applied to extract frequencies between 0.01-0.1 Hz, representatives of the low rs-fMRI Blood Oxygen Level Dependent (BOLD) signal. The signal from ventricles was regressed out using a least square approach in order to reduce non-neural correlations from the cerebrospinal fluid.

Several analysis approaches were applied on this data as following:

##### Graph theory-based analysis of functional connectivity

To evaluate direct correlations between spatially separated regions of interest (ROIs), Partial Correlation (PC) analysis was performed (Hübner et al., 2017) between the mean time courses of the BOLD rs-fMRI signal of the included ROIS. 37 ROIs covering the major mouse brain cortical and subcortical areas were extracted from the Allen Mouse Brain Atlas (AMBA) (2011 Allen Institute for Brain Science, Allen Mouse Brain Atlas available from http://mouse.brain-map.org/static/atlas) (Lein et al., 2007) and registered on morphological template using an in-house built MATLAB tool.

The following analysis steps were carried out:

i. Groups specific average FC matrices of 37×37 ROIs (corresponding to each experimental group: C57BL/6N and BALB/cJ mice) were generated and the matrices were arranged according to the most similar pattern of connectivity across ROIs. The matrices were Fisher transformed to obtain the z scores associated to the connection strength – the edges - between pairs of ROIs. A one-sample t-test (p<0.05, FDR corrected) was applied to identify the statistically significant pairs of connections (edges). The resulting significant FC matrices were used to as inputs for graph theory measures. For each group, the most influent nodes of the network were identified using as parameters the weighted hubness (presented as the size of the node in the graph) and the Stouffer coefficients (color coded).
ii. Direct intergroup (C57BL/6N vs. BALB/cJ) statistical comparison of connectivity matrices (P<0.05, uncorrected) was performed to identify the most different “nodes” and “edges”. A group comparison matrix (GCM) was generated and graphically represented (e.g. Figure 4G) with a color-scale, displaying statistically significant inter-group differences of connectivity. A method to identify most different network nodes among the FC matrices of the two strains was implemented. The following information was taken in account for ranking the most different nodes: the number of significantly changed connections for each node, the strength of the connection difference between pair of nodes and the number of statistically different connections of the neighboring nodes. This information is graphically represented in the GCM in relationship with the node’s size and ranked in Figure 4H. Additionally, the group comparison matrix was used for calculating the Stouffer coefficients (Stouffer et al., 1949) for each node. A single p-value was computed for each region based on the combination of the p-values derived from the statistical tests made on the correlations with all other regions. The results were color-coded for most significantly different nodes (e.g. nodes color in the Figure 4B, E, G).

**Seed-based functional connectivity analysis** was further performed with a MATLAB tool developed in-house. Several ROIs (extracted from AMBA) were selected for this analysis based on the DTI and VBM results, as well as based on the previously published reports (Anderzhanova et al., 2013; Fairless et al., 2012; Kim et al., 2012; Kumar et al., 2012; Ohl et al., 2001, p. 6; Sankoorikal et al., 2006; Wahlsten, 1974) on the behavior of C57BL/6N and BALB/cJ mice. These ROIs were:

i. Right and left somatosensory cortex (SSr and SSl) and right and left motor cortex (MOr and MOl). These ROIs were used to check the inter-hemispherical connectivity.
ii. Anterior cingulate area (ACA) and retrosplenial cortex (RSP), to derive information about the default mode patterns in the two mouse strains. c) Key players of the known reward aversion circuitry: accumbens nuclei (ACB), hippocampus (hc).

Partial correlation coefficients (Spearman correlation) were computed between the mean rs-fMRI time course of each ROI and the time courses from each voxel of the brain while excluding the effect of all the remaining voxels. Functional connectivity maps for each ROI were constructed at the individual level and also at the group level. Voxel-wise group based statistical analysis was performed using a two-sample t-test via SPM8 and the statistical results were cluster corrected at the P<0.05, FDR cluster corrected.

##### Directional connectivity analysis (DCA)

To probe dominant asymmetric information flow alterations within small network, population level directionality analysis using MVGC toolbox was performed (Barnett and Seth, 2014). A small network was constructed via selection of six regions of interest that showed structural/function strain differences in previous analysis: nucleus accumbens (ACB), anterior cingulate area (ACA), amygdala (AMY), caudate-putamen (CP), hippocampus (hc) and ventral tegmental area (VTA) (Figure 9 A). Briefly, the pre-processed resting state functional data was deconvolved (Wu et al., 2013) and pairwise unconditional Granger causality (completely data driven approach) (Barnett and Seth, 2014; Roebroeck et al., 2005) was carried-out on all possible seed pairs included in the constructed network, for each subject. Akaike information criterion (AIC) was engaged to find the best stable model order. Statistical analysis was implemented following t-test with multiple hypothesis testing (FDR correction, P<0.05).

## 3 Results

### 3.1 DTI reveals strain differences in structural connectivity

Diffusion tensor imaging data obtained from the BALB/cJ and C57BL/6N cohorts were used for global mouse brain fiber tracking, generation of hrFM as well as mapping of FA, RA, AD and trace values. These maps were assessed at individual and group levels. hrFM, displaying color-coded fiber orientations, were used as a first step to explore the whole brain structural connectome. Figure 1 presents exemplary hrFMs, representative for each strain (Fig.1-A for BALB/cJ strain and B for C57BL/6N strain). These maps are highlighting the variability of inter-hemispherical connectivity through corpus callosum (cc) within the BALB/cJ strain (Fig.1-A). Certain individuals from BALB/cJ group had visibly shorter cc; the variability along this major inter-hemispherical pathway is clearly identified at different callosal levels; namely the genu (gcc), body or middle part (mcc), and splenium (scc) parts (see Fig.1 and Video 1). C57BL/6N strain, in contrast, showed more homogenous and well-defined structure of this region (Fig.1-B).

**Figure 1.**
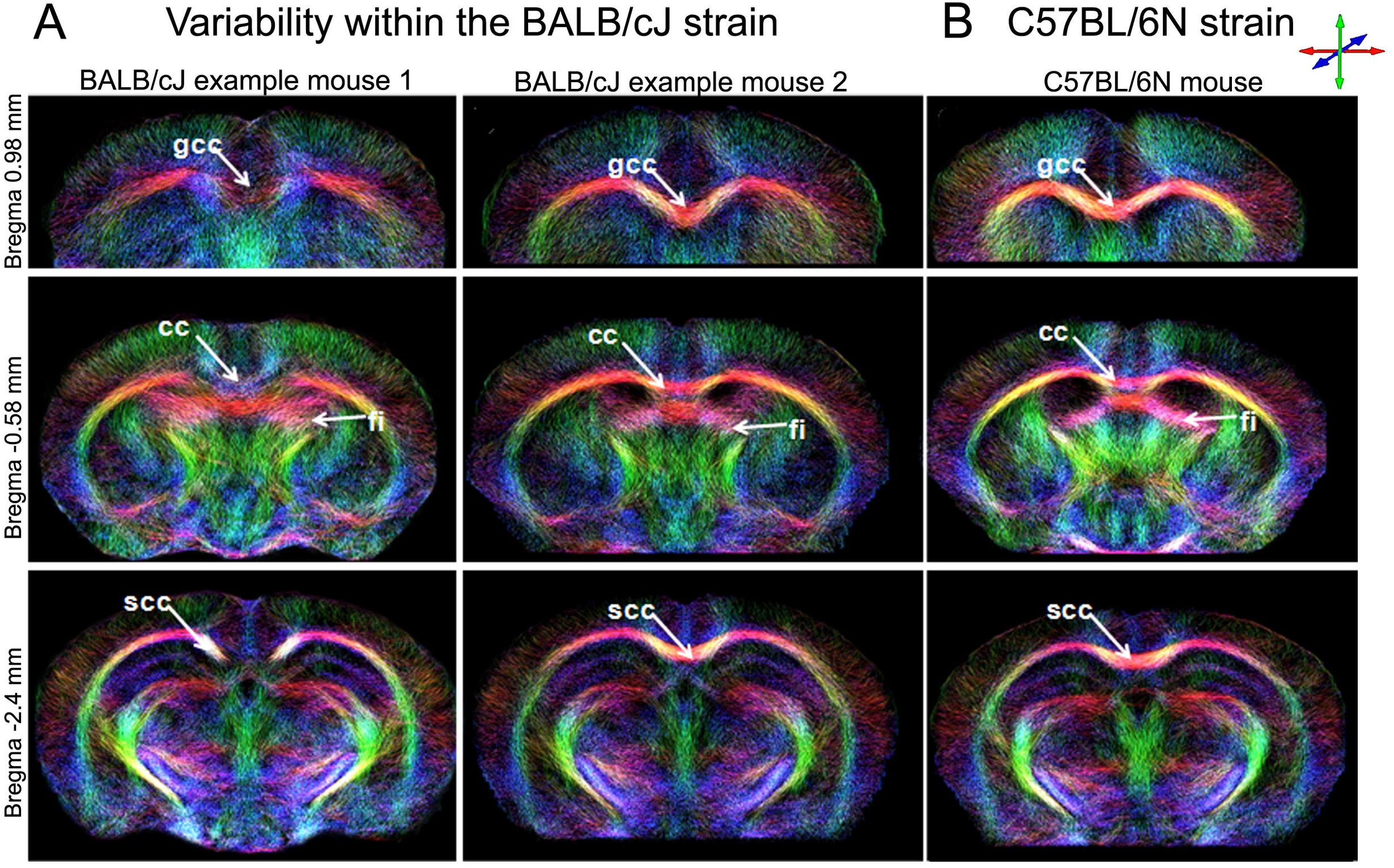
Living mouse brain connectional anatomy in BALB/cJ and C57BL/6N mouse brains. Representative high-resolution fiber maps (hrFM) were generated from the global fiber tracking data and were reconstructed with a resolution of 12×12×50 μm3, at different bregma levels (bregma −0.6; −0.7 and −0.9). **(A and B)**: Variability within the BALB/cJ mice population along the patterns of callosal (cc) connectivity. Reduced inter-hemispherical connectivity is seed in BALB/cJ mouse 1 compared to BALB/cJ mouse 2, at the levels of genu of the corpus callosum (gcc), body of the corpus callosum (bcc) and splenium of the corpus callosum (scc), suggestive of shorter callosal size in mouse 1. **(C)** Overall view of the connectional architecture in a C57BL/6N individual. The color-coding indicates the local fiber orientation: red, mediolateral; green, dorso-ventral; blue, rostro-caudal.

To further assess the inter-strain differences, group-specific mean FD maps were generated (Supplementary Figure 1-A and B) illustrating strain FD differences along the length of the cc. Individual FD maps were further used for voxel-wise statistical group comparison (C57BL/6N vs. BALB/cJ) (Suppl. Fig. 1-C and D). The C57BL/6N strain shows higher FD than BALB/cJ mice (p<0.05, FDR corrected; Suppl. Fig. 1-C) at the level of WM structures such as gcc and scc, fasciculus retroflexus (fr) or cerebral peduncle (cp). Significant differences are demonstrated in GM regions including prelimbic and anterior cingulate areas (PL and ACA), caudate putamen (CP) and dorsal hippocampus (dhc), temporal association areas (TEa), thalamus (TH), periaqueductal gray (PAG), and midbrain nuclei (MB). The shape of the areas is suggestive of FD differences along striato-cortical pathways (scp), reaching notably the ACA. By contrast, the statistics indicate higher FD values in BALB/cJ strain compared to C57BL/6N (Suppl. Fig. 1-D, p<0.05, FDR corrected) in motor areas (MO). The lateral ventricle (LV) area is marked in the statistics (Suppl. Fig. 1-D) as a result of morphological differences at the ventricular level in-between two strains, which is taken into account in the VBQ statistical method (Draganski et al., 2011).

Statistical analysis of FA maps at the group level showed comparable differences between strains: cc, including anterior forceps (fa), cingulum (cg), gcc and scc, internal capsule (int), medial forebrain bundle (mfb) and cp as well as gray matter or mixed gray/white matter regions: PL/ACA, TEa, CP, dhc, TH, midbrain and pontine reticular nuclei (MRN; PRN, respectively) demonstrated significantly higher FA values in C57BL/6N strain (p<0.05, FDR corrected, Fig. 2-A). The patterns are also indicative of increased FA along striato-cortical pathways. BALB/cJ mice, on the other hand, showed higher FA in ventral CP, as well as along caudal thalamo-cortical pathways (tcp), reaching somatosensory (SS) and auditory cortices (AUD) (p<0.05, FDR corrected, (Fig. 2-B).

**Figure 2.**
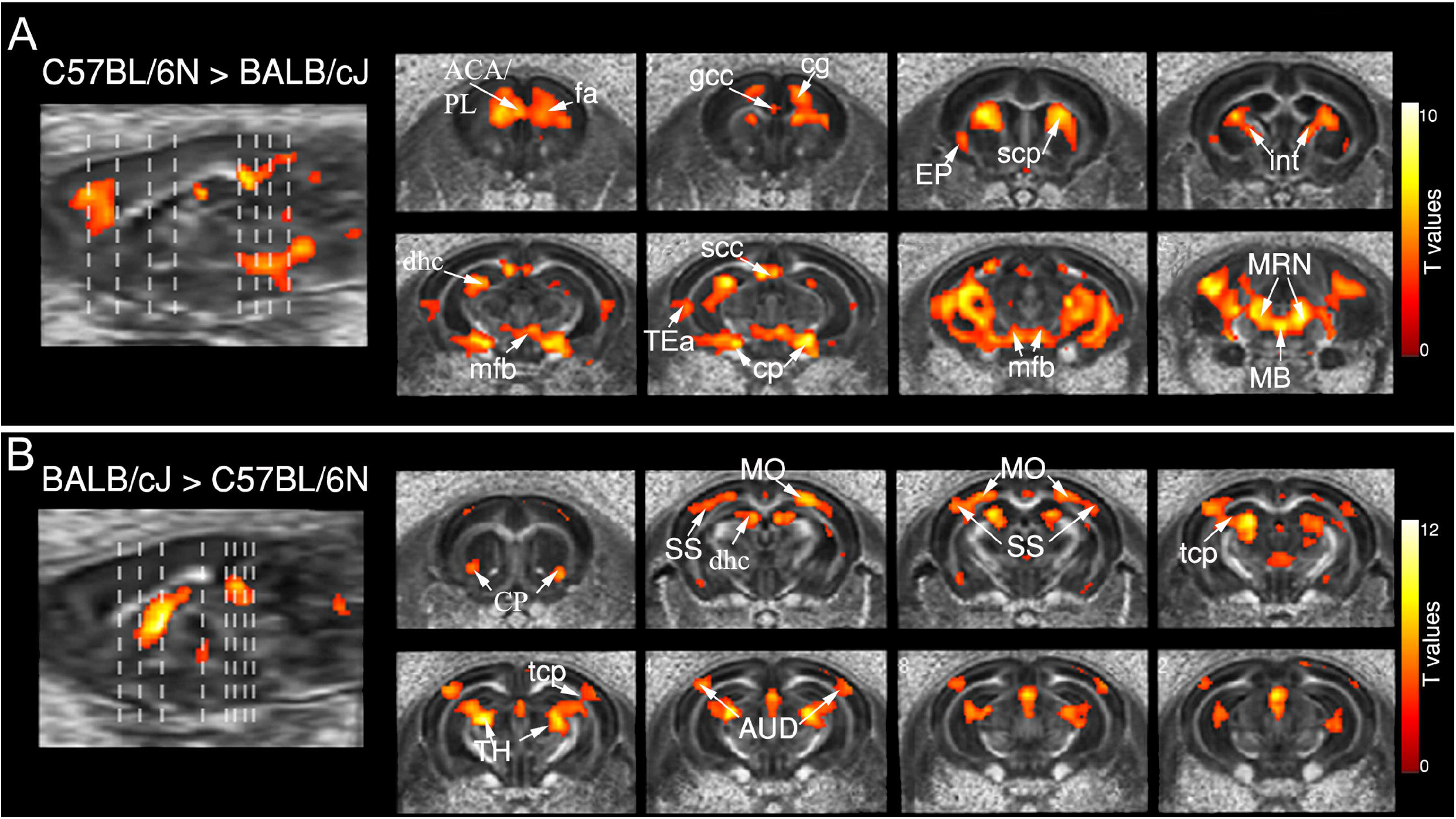
Fractional anisotropy (FA) maps reveal differences between strains. **(A-B)** Voxel-wise statistical group comparison of FA maps (p< 0.05, after false discovery rate (FDR) p-value correction; color scales represent t-values) indicate areas of differences **(C)** Contrast C57BL/6N > BALB/cJ: higher FA in C57BL/6N mice vs. BALB/cJ group. **(D)** Contrast BALB/cJ > C57BL/6N: higher FA in BALB/cJ mice vs. C57BL/6N group. [Abbreviations: Corpus callosum (cc), anterior forceps (fa), genu and splenium of cc (gcc and scc), internal capsule (int), medial forebrain bundle (mfb), cerebral peduncle (cp), prelimbic area (PL), caudate-putamen (CP), Ammon’s horn (CA), temporal association areas (TeA), thalamus (TH), midbrain and pontine reticular nuclei (MRN; PRN, respectively) dentate gyrus (DG), thalamo-cortical pathways (tcp), somatosensory (SS) and auditory cortices (AUD).]

### 3.2 Inter-strain regional volumetric differences uncovered by VBM

Figure 3 shows statistically significant brain volume differences in-between strains. C57BL/6N mice had significantly bigger (p<0.05, FDR corrected; Fig. 3-A) frontal cortical areas, including ACA, MO and SS as well as TEa and entorhinal areas (ENT). At subcortical levels, the analysis revealed bigger septal areas in C57BL/6N mice and increased size at the level of thalamic nuclei (TH) (i.e. ventral group of the dorsal thalamus). The results also indicate larger lateral ventricles (LV) in the C57BL/6N strain. For BALB/cJ mice, volumes were significantly bigger (p<0.05, FDR corrected; Fig. 3-B) in piriform area (PIR), CP, certain hippocampal areas, medial hypothalamus (HY), substantia nigra (SN), VTA, and PAG as well as the middle part of the cc.

**Figure 3.**
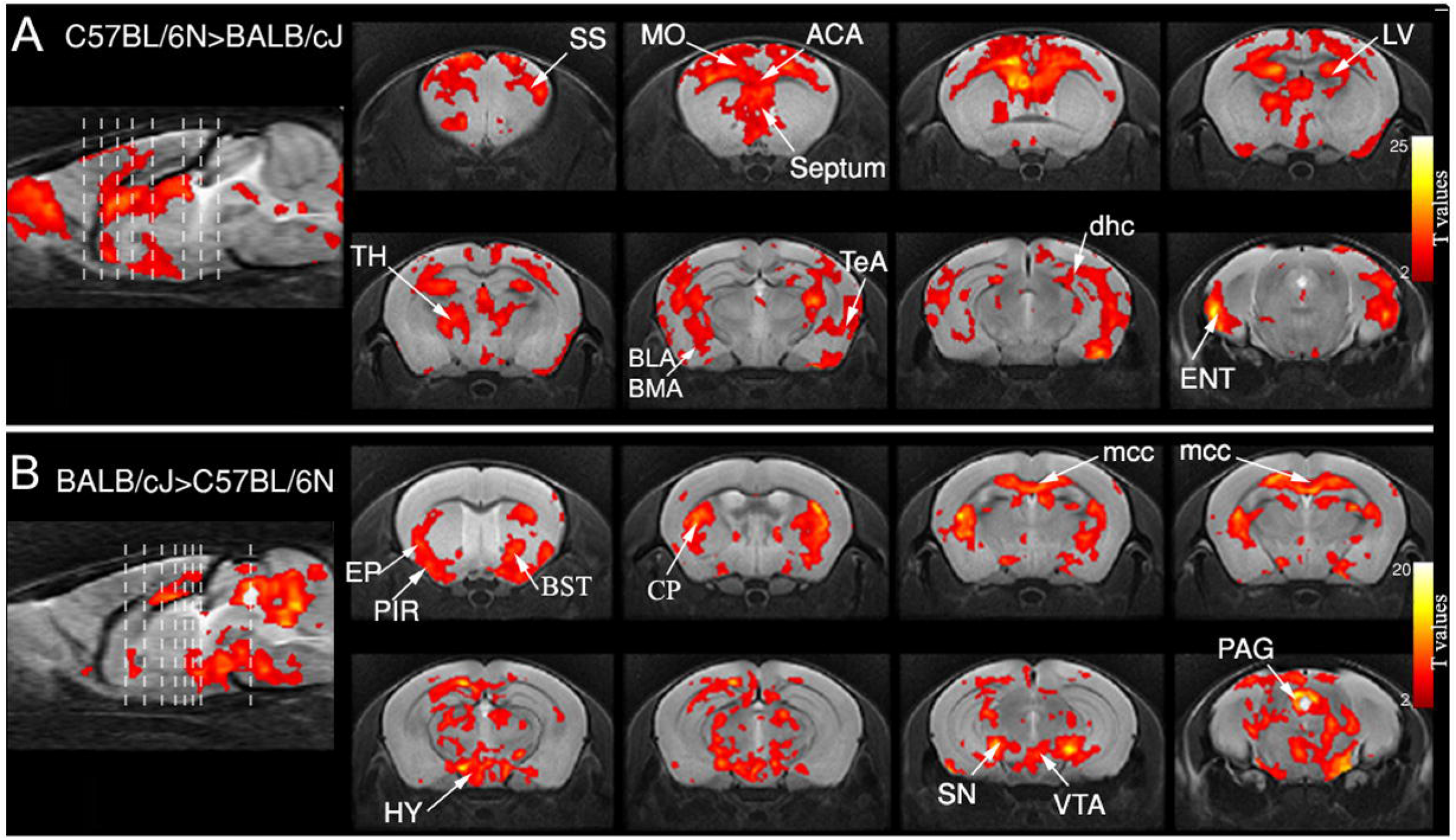
Inter-group statistical comparison for voxel-based morphometry (VBM). (Threshold at P< 0.05 after false discovery rate (FDR) p-value correction; color scales represent t-values). **(A)** Contrast C57BL/6N > BALB/cJ. **(B)** Contrast BALB/cJ > C57BL/6N. [Abbreviations: Anterior cingulate area (ACA), motor areas (MO), sensory areas (SS), temporal association areas (TeA), entorhinal areas (ENT), thalamus (TH), lateral ventricles (LV), piriform area (PIR), caudate-putamen (CP), Ammon’s Horn (CA), hypothalamus (HY), substantia nigra (SN), ventral tegmental area (VTA), periaqueductal gray (PAG), corpus callosum (cc).]

### 3.3 Functional connectivity patterns in C57BL/6N and BALB/cJ mice

To complete the picture of the brain connectome in the two investigated strains we further mapped the large-scale FC patterns via rs-fMRI, to determine whether different brain structural architectures are accompanied by functional discrepancies.

#### 3.3.1 Global FC features of C57BL/6N and BALB/6J brains

We applied graph network analysis at group level and mapped the topological organization of the FC, by modeling the 37×37 ROI PC matrices (Fig. 4-A and D). The nodes were defined by the 37 ROIs extracted from AMBA, covering major cortical and subcortical brain areas. The selection of nodes was also guided by the structural results. For instance, certain nuclei, such as EP were included because differences in this area were observed in the voxel-wise FA analysis. Prominent inter-strain differences were obtained in the “hubness” characteristics (Fig. 4-B, C vs. E, F). In the C57BL/6N group a dominance of basal forebrain subcortical areas was revealed (Fig. 4-C – blue labels), encompassing the major limbic centers modulating reward (ACB, CP, ventral pallidum-PALv), stress and anxiety (bed nuclei of stria terminalis - BST) but also integrative centers for sensory information (Claustrum – CLA and EP). Cortical hubs included frontal areas; namely, insula (AI), piriform (PIR) cortex and orbito-frontal cortex (ORB), but also primary motor (MOp). The key player of the default mode network - the retrosplenial cortex (RSP) – was one of the C57BL/6N FC hubs.

**Figure 4.**
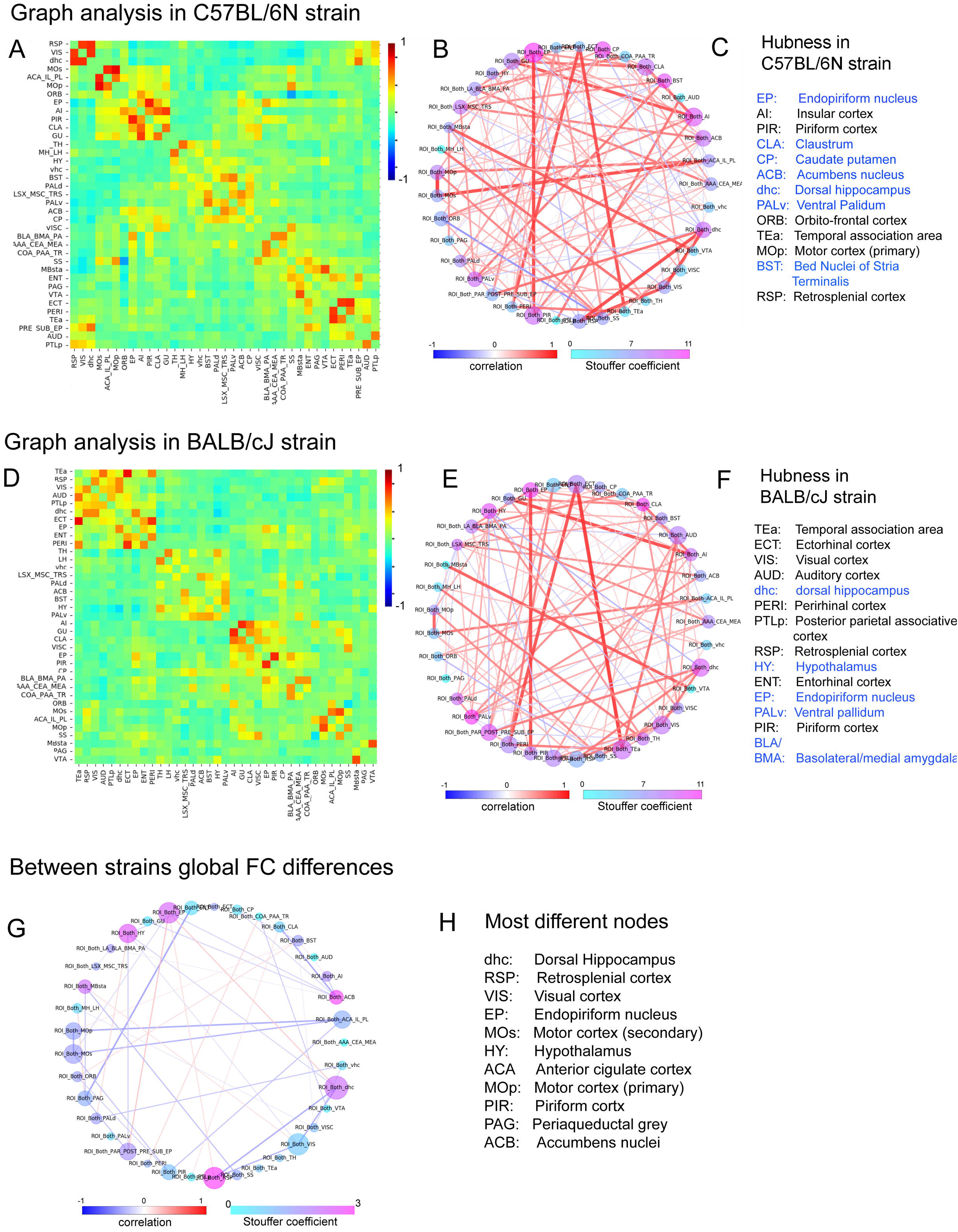
Functional connectivity matrices and related graph theoretical measures for C57BL/6N and BALB/cJ mice. (A and D) Functional connectivity matrices created with 37 bilateral brain areas (nodes) are represented for **(A)** C57BL/6N and **(D)** BALB/cJ. Color scale denotes correlation coefficients. **(B and E)** Statistically significant connections presented on a scale of correlation coefficients and matrix nodes color coded on a scale of Stouffer coefficients (one sample t-test, p<0.05, FDR corrected) for **(B)** C57BL/6N and **(E)** BALB/cJ matrices. (Node sizes correspond to hubness ranking.) **(C)** C57BL/6N and **(F)** BALB/cJ brains have distinct hub regions (List is ordered according to ranking, highest to lowest). **(G)** Differences between C57BL/6N and BALB/cJ brain FC are shown (Two sample t-test, p<0.05, uncorrected.). Connection strength differences are depicted on a scale of correlation coefficients and nodes are color coded to depict differences according to Stouffer coefficients; node sizes correspond to most different areas. **(H)** Ranking of most different nodes between C57BL/6N and BALB/cJ.

The “hubness” patterns in the BALB/6J mice was dominated by cortical areas (see Fig. 4-F), including associative (TEa) and sensory areas (visual - VIS, auditory - AUD, PIR). A marked difference when compared to the C57BL/6N was the absence of striatal/reward-related regions among the identified relay centers of the FC (i.e. ACB, CP).

Direct inter-group comparison of the global FC matrices identified the brain areas with divergent patterns of connectivity (Fig. 4-G, H). ACB showed highest number of different connections (see color coding – magenta). However, when normalized for the strength of the connectivity difference with other nodes, dorsal hippocampus emerged as the most dissimilar area in terms of FC (Fig. 4-H – and 4-G – size of the nodes). Furthermore, our global analysis suggested strain variations in the DMN features, as two of the main DMN nodes (ACA and RSP) appeared among the nodes with variant FC (Fig. 4-H). Several sensory processing areas (VIS, PIR, EP) as well as MO suggest also strain specific connectivity patterns.

Most of the areas indicated as divergent in the graph resting state network analysis (Fig. 4-G and H) were also highlighted in the inter-strain comparison of the structural data (i.e FA voxel vise analysis from Fig.2). Therefore, the next group level seed analysis of FC was guided both by the structural results and the brain-wide graph theoretical resting state results.

#### 3.3.2 Inter-hemispherical FC

Given the structural differences observed at callosal level in between the two strains, we first evaluated the inter-hemispherical FC, largely mediated through cc connections. The connectivity between the right and left hemispheric SS and MO cortical areas were therefore investigated. We selected successively right and left-lateral SS (SSr and SSl) as seed regions for mapping their connectivity patterns across the whole brain; subsequently, group comparisons were performed (two-sample t-tests, p<0.05, FDR cluster corrected). Despite the group differences observed at the level of gcc and scc after group comparison for structural FD and FA maps, the FC of the SS cortices with the same anatomical areas of the contralateral hemisphere (homotopic areas) does not show statistically significant inter-group differences (Fig. 5-A and B). Interestingly, the results indicate greater synchrony of the BOLD signal ipsilaterally, within the cortical areas of the same hemisphere for the BALB/cJ mice (Fig. 5-A and B, blue areas). The feature was reproducible for both right and left SS seeds. SS cortex also showed stronger correlation with RSP and VIS area in C57BL/6N strain (Fig. 5-A, red areas).

**Figure 5.**
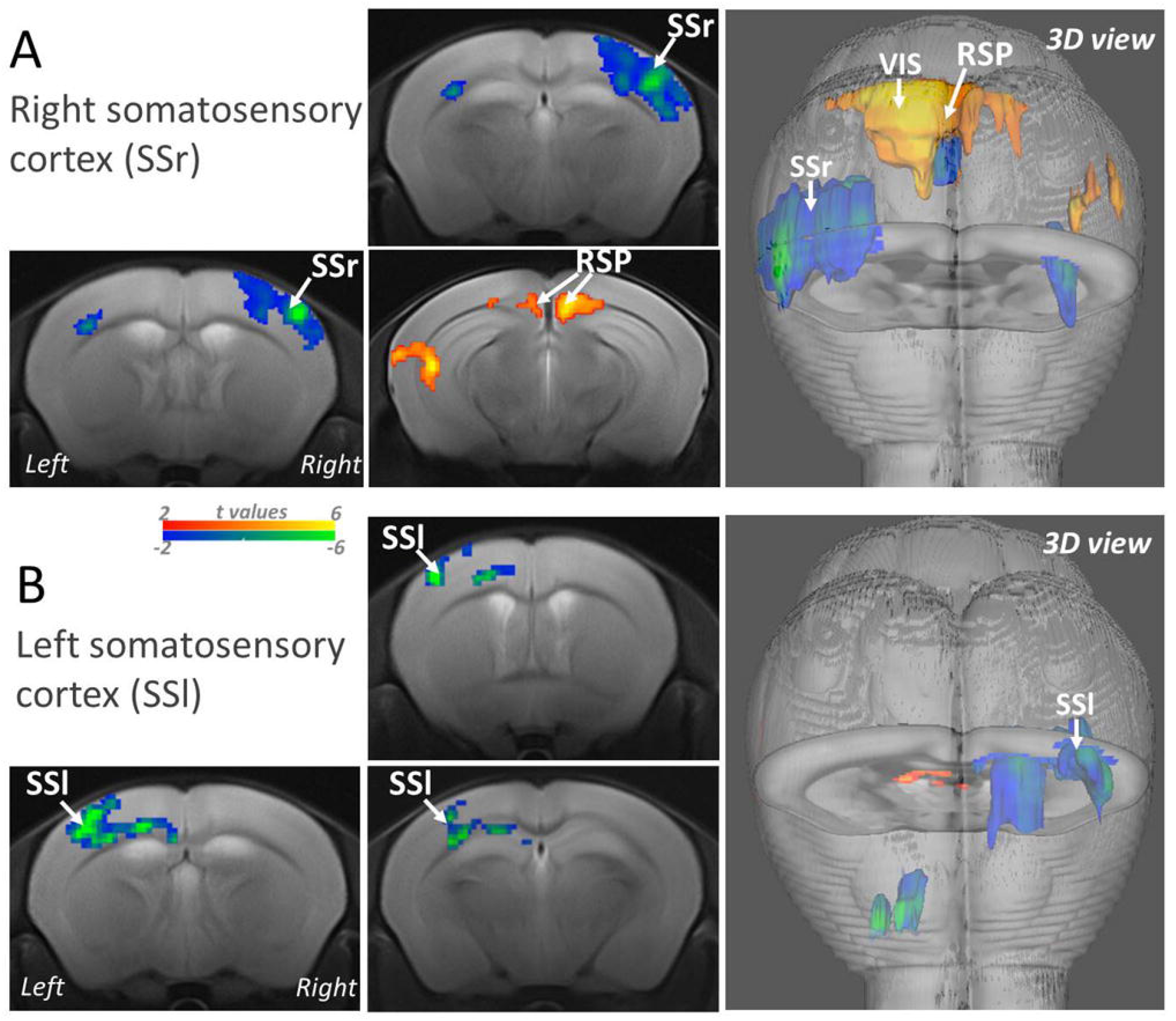
Inter-group differences in somatosensory cortex functional connectivity for C57BL/6N and BALB/cJ. Inter-group statistical comparison results of seed-based connectivity maps for **(A)** Right SS; **(B)** Left SS regions of interest (ROI). (FWER correction applied at cluster level for p<0.05); Scales represent t-values for contrast C57BL/6N > BALB/cJ in red or BALB/cJ > C57BL/6N in blue). [Abbreviations: somatosensory area right and left (SSr and SSl); posterior parietal association areas (PTLp), retrosplenial cortex (RSP), visual area (VIS).]

Inter-group statistical analysis carried-out for MO areas - right (MOr) - and left (MOl), indicated more complex differences of FC. First reproducible feature for both right and left MO was the stronger FC connectivity with frontal cortical areas (ACA but also within rostral MO – ipsilateral and contralateral to the seed) and frontal subcortical areas (including ACB, CP, BST) in C57BL/6N. However - along the rostro-medial-caudal axis, the medial brain regions - MO (right and left) displayed higher intra- and inter-hemispherical synchrony of the BOLD resting state signal in the BALB/cJ group (Fig. 6, blue patterns), notably within the sensory-motor cortex (SS and MO).

**Figure 6.**
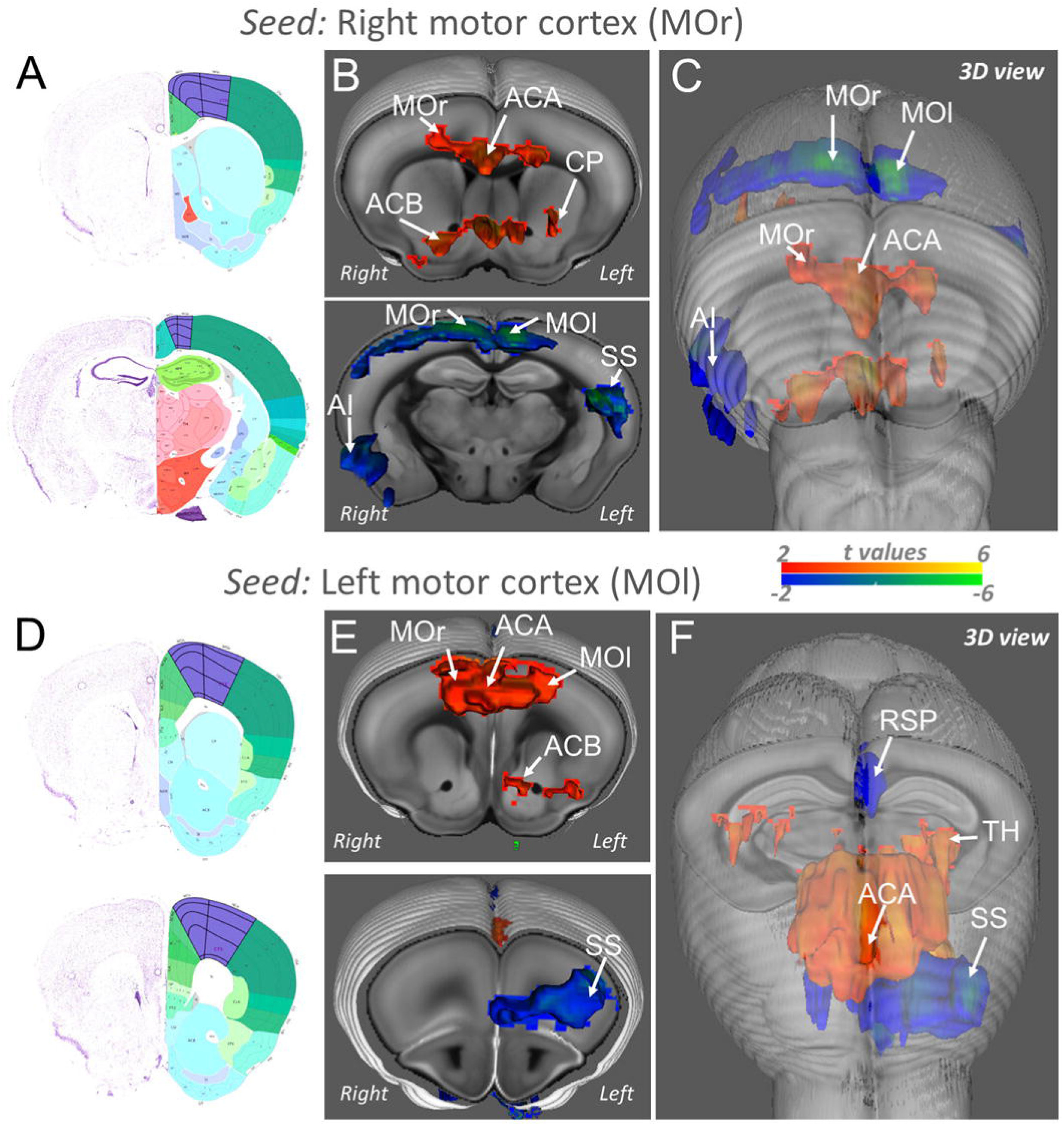
Inter-group differences in motor cortex functional connectivity for C57BL/6N and BALB/cJ. Inter-group statistical comparison results of seed-based connectivity maps for **(B, C))** Right motor cortex (MOr); **(E, F)** Left motor cortex (MOl) regions of interest (ROI). (FWER correction applied at cluster level for p<0.05; Scales represent t-values for contrast C57BL/6N > BALB/cJ in red or BALB/cJ > C57BL/6N in blue). **(A, D**) The atlas slides (left panel: from Allen Mouse Brain Atlas) corresponding to the presented MRI images indicate in magenta the motor areas. [Abbreviations: motor area right and left (MOr and MOl), anterior cingulate area (ACA), caudate-putamen (CP), nucleus accumbens (ACB), agranular insula (AI), somatosensory area (SS), retrosplenial area (RSP).]

#### 3.3.3 DMN patterns in C57BL/6N and BALB/cJ strains

The significant strain-specific features measured with structural MRI (Figs. 2 and 3) in the frontal brain areas including ACA - that is part of DMN- and along frontal part of cg bundle (Fig. 2-A) suggested that DMN patterns might also have strain-specific characteristics. Therefore, we probed the topological patterns of DMN using seed analysis. ACA and RSP were previously described as the key DMN nodes in rodents, showing synchronous rs-fMRI activity. We used the RSP as seed and mapped the spatial extent of DMN network, shown comparatively in Figure 7 for the two strains. A prominent pattern of BOLD rs-fMRI signal synchrony is noted along top cortical areas in both groups, encompassing the RSP and ACA cortices. Two main features are relevant: (i) larger cortical extent of the BALB/cJ DMN, along the caudal to rostral axis and (ii) extension of the RSP connectivity patterns towards subcortical areas (CP, hc and TH) in C57BL/6N mice. Inter-group comparative evaluation (Fig. 7-B, p<0.05, FDR cluster corrected) confirmed significantly stronger connectivity of RSP towards dhc and TH areas in C57BL/6N group. Further inter-group analysis of ACA FC (Fig. 7-C) unmasked major differences in the topology of ACA network: (i) stronger patterns of synchrony within rostral ACA and with neighboring cortical areas (MO) in C57Bl/6N mice and (ii) stronger ACA connectivity with RSP and dhc in the BALB/cJ group.

**Figure 7.**
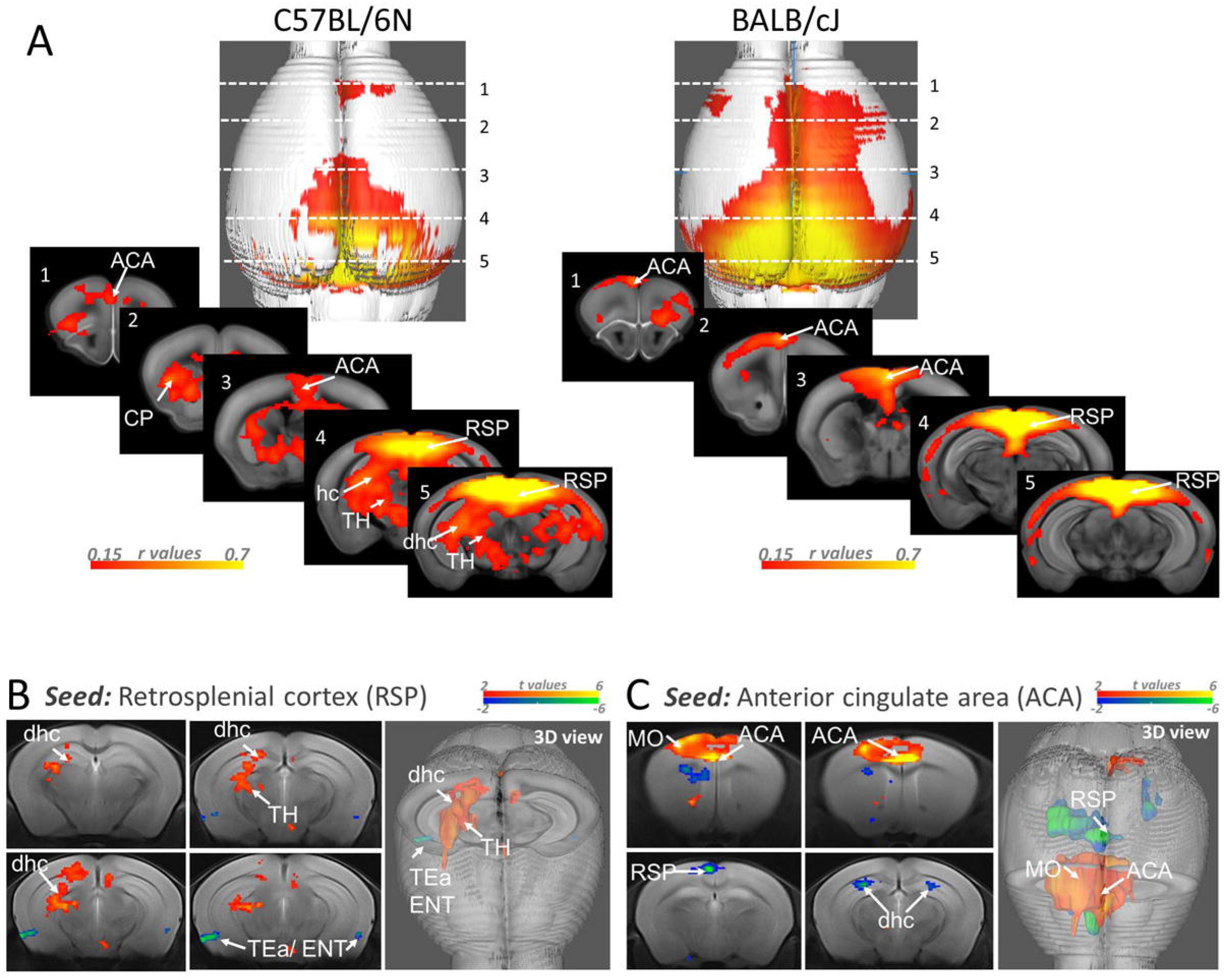
Default mode network (DMN)-like patterns in C57BL/6N and BALB/cJ. **(A)** Default mode network as mapped using retrosplenial area (RSP) as seed in the C57BL/6N and BALB/cJ strains. (Correlation values were scaled between 0.15 and 0.7.) **(B and C)** Inter-group statistical comparison of functional connectivity in (B) Retrosplenial area (RSP) and (C) Anterior cingulate area (ACA); (FWER correction applied at cluster level for p<0.05; Scales represent t-values for contrast C57BL/6N > BALB/cJ in red or BALB/cJ > C57BL/6N in blue) [Abbreviations: motor area (MO), dorsal hippocampus (dhc), thalamus (TH), temporal association area (TEa), entorhinal area (ENT).]

#### 3.3.4 Functional connectivity of the reward/aversion centers

We next performed fine-grain connectivity mapping of ACB and hc using seed analysis and carried-out inter-group statistical analysis (Fig. 8-A and B). These two areas are among key nodes of the limbic and/or reward-aversion system. This choice was motivated by reported behavioral results in the two strains, indicating different sociability and anxiety-related behaviors, possibly reflecting strain-specific patterns of the limbic FC. FC modifications of the limbic subcortical areas may emerge as well on the basis of the structural differences observed in the striatum (i.e greater FA and FD along C57BL/6N striatal-cortical pathways – Fig. 2-A) or dhc (Fig. 2-A and B), as shown in our study. Additionally, graph-based network analysis identified dhc and ACB as major nodes with divergent FC features in the two experimental groups (Fig. 4-G and H). Seed correlation approach – in agreement with FA increase along striato-cortical pathways in C57BL/6N mice - revealed better rs-fMRI signal synchrony between ACB and frontal cortex, encompassing ORB, ACA and MO in this strain (Fig. 8-A), but also more caudally with RSP. Subcortically, ACB showed locally stronger FC with other centers of the reward circuitry, notably CP and ventral tegmental areas VTA (Fig. 8-A) for C57BL/6N strain. Seed-based cartography of the hc network showed a better synchrony of the hc rs-fMRI signal with medial ACA, AI, SS, TEa as well as RSP in C57BL/6N brains, while better correlating with TH nuclei in BALB/cJ strain (Fig. 8-B).

**Figure 8.**
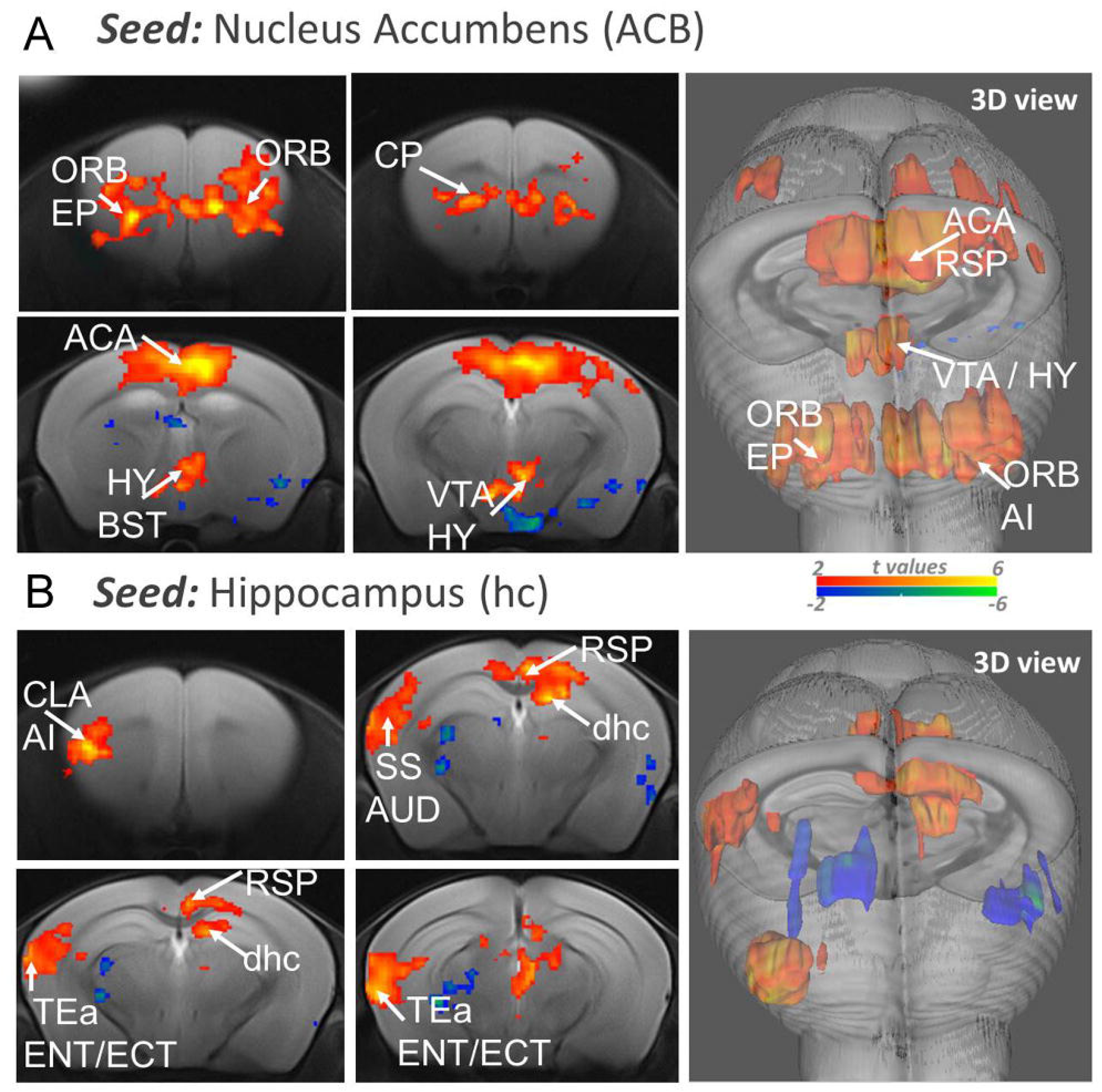
Inter-group statistical comparison of seed-based connectivity maps for nucleus accumbens and hippocampus. **(A)** Nucleus accumbens (ACB) and **(B)** Hippocampus (hc) functional connectivity differences between C57BL/6N and BALAB/cJ. (FWER correction applied at cluster level for p<0.05; Scales represent t-values for contrast C57BL/6N > BALB/cJ in red or BALB/cJ > C57BL/6N in blue) [Abbreviations: orbital area (ORB), anterior cingulate(ACA), endopiriform nucleus (EP), caudate-puatamen (CP), hypothalamus (HY), bed nucleus of stria terminalis (BST), ventral tegmental area (VTA), retrosplenial area (RSP), agranular insula (AI), claustrum (CLA), somatosensory area (SS), auditory area (AUD), temporal association area (TEa), entorhinal (ENT) and ectorhinal areas (ECT), dorsal hippocampus (dhc).]

#### 3.3.5 Directional communication analysis among key nodes of the reward/aversion network

To investigate the direction of information flow between pairs of nodes of the reward-aversion circuitry we conducted pairwise Granger Causality analysis. We constructed a network by selecting six relevant regions showing group differences in structural or seed-based FC analysis: ACB, hc, CP, ACA, amygdala (AMY) and VTA (Fig. 9-A). These regions are core players of the reward circuitry and known to be involved in regulating-among others-the affective/social behaviors. For each set of ROIs, we computed pairwise Granger Causality extracting bidirectional information between pairs of nodes (Fig. 9-B and D) and established dominant directionality (p<0.05; FDR corrected). Overall, the information flow directionality was similar in both groups with the exception of the ACA. This node was found to be dominantly receiving information from hc, AMY and CP only in C57BL/6N group and sending information towards ACB only in BALB/cJ strain; clearly providing evidence of group directional information differences in ACA region (Fig. 9-C and E). Indeed, these findings again highlight the strain-specific communication patterns of ACA region, in accordance with seed-based FC analysis and microstructural and morphological (DTI and VBM) results.

**Figure 9.**
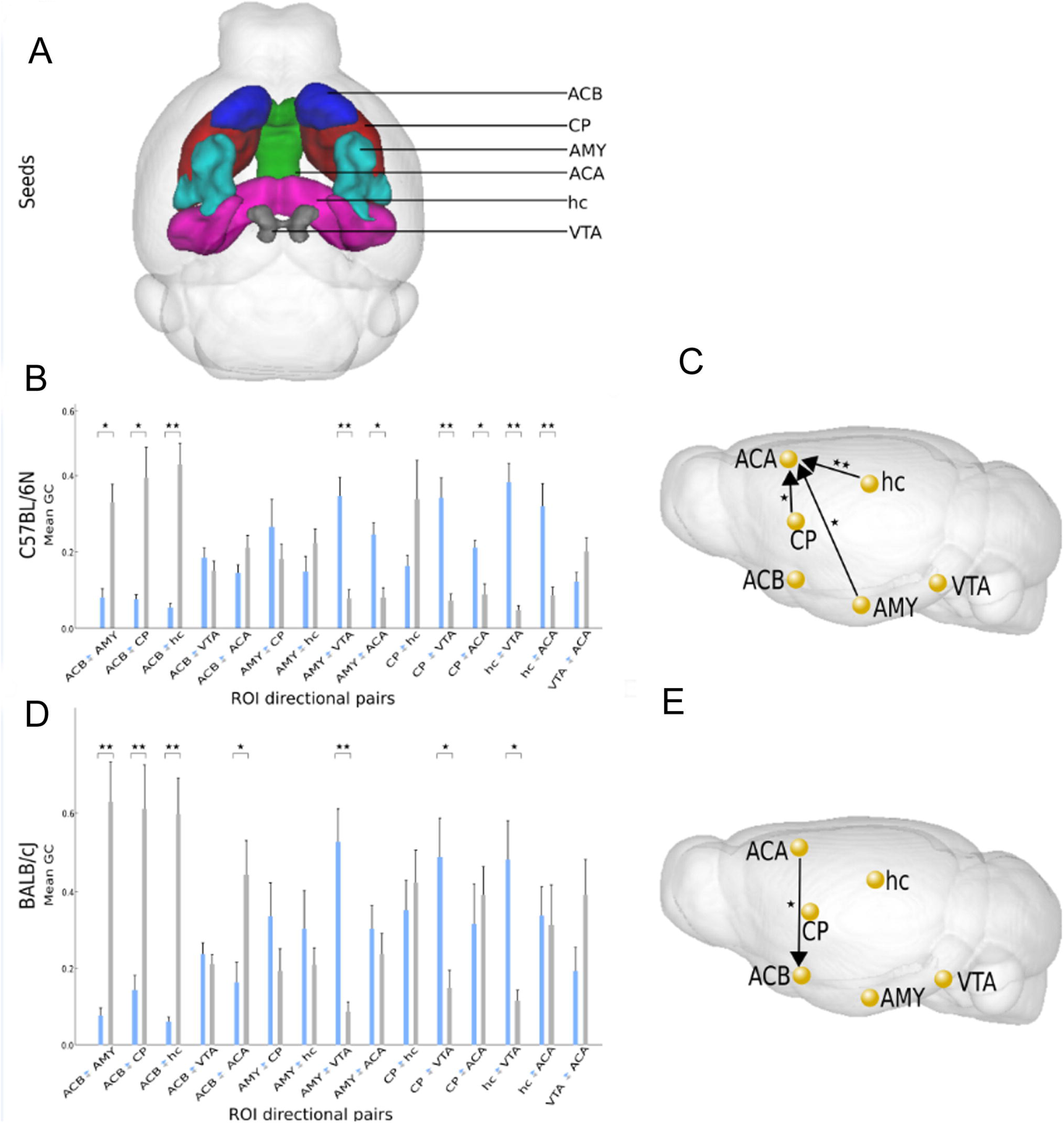
Directional connectivity between reward/aversion centers. **(A)** 3D individual color-coded representation of 6 selected seeds over mouse brain template with identification; **(B,D)** Mean (+SEM) of Granger Causality results with t-test (*p<0.05,**p<0.01,***p<0.001, FDR corrected). **(C,E)** Graphical illustration of dominant directionality difference between the two strains (C57BL/6N and BALB/cJ) with black directional arrows including black stars to represent level of significance. [Abbreviations: Nucleus Accumbens (ACB), anterior cingulate area (ACA), Hippocampus (hc), Amygdala (AMY), Caudate-putamen (CP) and ventral tegmental area (VTA)]

## 4 Discussion

In the recent years, huge effort has been dedicated for the characterization of the mouse brain connectome - at micro and mesoscale (Bardella et al., 2016; Grandjean et al., 2017; Grange, 2018; Ingalhalikar et al., 2014; Knox et al., 2018; Oh et al., 2014; Pervolaraki et al., 2019), as this species remains the principal animal model used in neuroscience research. Most of these studies were carried-out in the C57BL/6 strain, which remains the primary “genetic background” for modelling of human disease. However, C57BL/6 has many behavioral characteristics that make it useful for some work and inappropriate for others (Clipperton-Allen et al., 2015; Fairless et al., 2013; Fontaine and Davis, 2016; Ohl et al., 2001; Pilz et al., 2015; Sankoorikal et al., 2006; Yoshida et al., 2016). Therefore, the uncovering the large-scale brain circuitry configurations in a strain-specific manner represents a first step towards a better understanding of modifications in brain networks under the influence of various genetic factors, pharmacological interventions and pathological conditions. In this study, we systematically characterized the brain morphological patterns along with the structural and functional connectivity profiles of two commonly used mouse strains, C57BL/6N and BALB/cJ, via non-invasive *in vivo* MRI techniques. We show brain-wide morphological and functional differences encompassing cortical and subcortical structures as well as WM tracts. We further provide exemplary high-resolution fiber tractography maps demonstrating the inter-individual variability across inter-hemispherical callosal pathways in the BALB/cJ strain.

### 4.1 Strain specific morphological and structural brain connectivity patterns

Multiple previous histological and MRI studies captured the impact of genetic variability on the morphology of the brain (Fairless et al., 2012, 2008; Fontaine and Davis, 2016; Kim et al., 2012; Kumar et al., 2012; Zhang et al., 2015). Mice on different genetic backgrounds have stable yet distinct behavioral phenotypes that may lead to unique gene-strain interactions on brain structure. One classical example is observed with Fragile X Mental Retardation 1 knock-out (FMR1-KO) mice in which Fragile X Syndrome and related phenotypic manifestations of autism spectrum disorder (ASD) are induced. Indeed, while FMR1-KO in C57BL/6 strain show very few differences in brain morphology compared to wild-type mice (Ellegood et al., 2010) FMR1-KO mice on an FVB background displayed multiple neuroanatomical differences, including modifications of major white matter structures throughout the brain and changes in areas associated with fronto-striatal circuitry (Lai et al., 2016). White matter changes, including prominent reduction of callosal fibers are also reported with BTBR mouse, another ASD mouse model (Dodero et al., 2013; Ellegood et al., 2013; McFarlane et al., 2008; Miller et al., 2013) In our study, we produced highly resolved fiber maps that depict striking inter-individual differences along callosal commissure in the BALB/cJ female mice, with significant inter-hemispherical under-connectivity in the rostral and caudal cc in some individuals. These *in vivo* brain tractograms are in agreement with previous histological (Fairless et al., 2012; Moy et al., 2007; Wahlsten, 1974) and diffusion MRI findings in BALB/cJ males (Kim et al., 2012; Kumar et al., 2012) that reported variability in the size of cc, including the complete absence, in 30- 40% of this strain (Wahlsten, 1974). This variability was suggested to be possibly due to a variable delay in formation of an interhemispheric bridge of tissue in the dorsal septal region during prenatal development (Wahlsten, 1974). In some ASD relevant rodent models, this callosal underconnectivity normalizes overtime (Frazier et al., 2012), especially in the rostral regions of the cc - suggesting developmental trajectories that might influence behavioral outcome overtime. The BALB/cJ inbred strain was long been discussed as relevant for certain aspects of ASD (Brodkin, 2007), showing low sociability, high anxiety and aggressive behaviors. However, compared to C57BL/6J mice, only juvenile BALB/6J animals demonstrated lower sociability scores (Fairless et al., 2012) that positively correlated with mean diffusivity values along the external capsule of the WM (Kumar et al., 2012).

Although in our study, we cannot establish a direct link between behavioral features and strain-specific brain structural modifications, we provide a refined, high resolution evidence about fiber density and tissue anisotropy dissimilarities at the level of callosal structures (gcc, mcc and scc) and alongside other WM bundles (fi, fx, cg) in female C57BL/6N and BALB/cJ mice. GM regions, likewise, showed distinct microstructural patterns in the two mice populations, prominent differences appearing in frontal cortices (ORB, IL, PL, ACA); along the fronto-striatal pathways (within CP) and within thalamic and caudal midbrain nuclei, including dopaminergic VTA and substantia nigra areas. For all these areas BALB/cJ mice show lower fiber densities and FA. Changes in fronto-striatal circuitry have been often implicated in the fragile X syndrome (FXS) with autistic-like features (Dennis and Thompson, 2013a, 2013b) as well as other brain disorders (Qiu et al., 2011; Shepherd, 2013), including attention deficit disorders. In FXS patients, the lack of response inhibition and conscious regulation of anxiety are among the phenotypes that relate to the functions of these regions (Bonelli and Cummings, 2007; Eagle et al., 2008; Fox et al., 2010), so it might be relevant that fronto-striatal fiber pathways are less dense in BALB/cJ animals, that are known to show high anxiety levels.

Furthermore, we observed large volumetric variations between two strains in the prefrontal, rostral MO, SS and temporal association cortices along with septal, hc and TH areas displaying bigger volumes in C57BL/6N. Intriguingly, mcc region was found to be larger in BALB/cJ; perhaps compensating for shorter rostro-caudal cc length to ensure unperturbed inter-hemispherical information transfer. In an *ex-vivo* study comparing several mouse strains (Ellegood et al., 2015) for brain morphometry, BALB/cJ mice were found to have smaller frontal and parieto-temporal lobes, similar to our findings. Such regional brain volume changes were correlated with functional impairments in ASD patients. For example, changes in superior and medial prefrontal gyri volumes correlate with cognitive outcomes in spatial relations and verbal fluency scores in ASD adolescents (Bray et al., 2011). In addition, larger CP volumes– as seen in BALB/cJ mice - is a recurrent finding in ASD patients and are associated with repetitive behaviors and cognitive deficits (Peng et al., 2014).

### 4.2 From structure to function: divergent functional connectivity in C57BL/6N and BALB/cJ mice

To discover whether differences in the brain structural scaffolding of the two strains also give rise to functional variations, we first performed a global assessment of the topological features of the rs-fMRI FC in both strains via graph analysis. The “brain hubs” - a set of highly connected regions serving as integrators of distributed neuronal activity were defined for each strain. These FC nodes have an integrative role and are therefore susceptible points to dysfunction in brain disorders. Strain differences in cerebral network “hubness” may impose strain-specific regional dominance in processing of the functional information. The circuitry vulnerability to stressors might be divergent, as well as the circuitry response in terms of mal-adaptations or compensatory remodeling.

Based on the anatomical selections of the FC network nodes, our analysis revealed that the dominant players in the C57BL/6N strain were subcortical forebrain limbic areas, including endopiriform nucleus, claustrum, along with centers controlling reward and motivation (ACB, CP, dhc, PALv). The endopiriform nuclei as well as claustrum are intriguing brain structures, featuring the highest connectivity per regional volume in the brain. Their connectivity patterns were dissimilar, both in structural and functional analysis. Enclosed between the striatum and the insular cortex, with widespread reciprocal connections with the sensory modalities and prefrontal cortices, these nuclei seem to perform functions in processing limbic and sensorimotor information (Watson et al., 2017). The “hubness” in BALB/cJ strain is dominated by cortical areas with associative and sensory valence (Tea, VIS, AUD, PERI, PTLp, PIR) but also includes aversive centers, such as amygdala (BLA/BMA).

In both strains, the RSP seem to appear as a hub, emphasizing the importance of this DMN node in the FC of the mouse brain. Inter-group comparison highlighted FC differences in brain areas showing variations in either the FA or FD maps or the morphometry measures. Dhc, the most different node in terms of FC showed higher FA and FD values, but also bigger volumes in the C57BL/6N strain. This is coherent with previous studies (de Sá-Calçada et al., 2015) that found smaller dendritic lengths and fewer ramifications in dentate granular neurons of BALB/cJ hippocampus compared to C57BL/6 strain. Differences in the hippocampal functioning and contextual memory formation in between the two strains were among the earliest demonstrated features (Chen et al., 1996) as hippocampal lesions impact less on contextual fear conditioning in BALB/c mice than C57BL/6 animals. In another study hippocampus related cognitive deficits – poor learning and memory performance in both the open field and passive avoidance inhibitory tasks - were assessed in stressed BALB/c, but not C57Bl/6 mice (Palumbo et al., 2010). This feature can be also mediated by distinct FC between dhc and frontal cortex (including ACA) and more dominant flow of information passing from the dhc to the ACA in C57Bl/6N strain. Previous studies demonstrated a role for the ACA-dhc pathway in recall of remote memory and its involvement in attention in both novel situations as well as during performance of well-learned tasks (Weible, 2013).

We further examined if inter-hemispherical FC of homotopic cortical areas, known to be largely mediated by callosal commissure show strain-specific features. FC between SS and MO homotopic areas did not show clearly a stronger inter-hemispherical connectivity in the C57BL/6N mice, as one might expect on the basis of the structural findings. Rather, a very specific pattern of FC differences was observed, along the rostro-caudal axis. For SS cortex, ipsilateral connectivity was stronger in the BALB/cJ mice, displaying a greater intra-hemispherical synchrony of the BOLD rs-fMRI signal. These results could be reproduced for the MO cortex in the medial brain areas, where better inter-hemispherical FC was also noticed for BALB/cJ. These results reinforced the observed higher intra cortical FD and FA values of the motor cortex in the BALB/cJ strain and bigger volume of the cc in the medial region, eventually favoring the functional brain communication locally. The pattern is reversed however in more rostral regions where seed analysis of MO cortex shows stronger intra and inter-hemispherical FC in the C57BL/6N animals. Higher density of fibers and bigger brain volumes measured in the C57BL/6N strain for the ACA, ORB regions and along gcc, might facilitate the FC in this strain locally.

In a study comparing inter-hemispherical FC in different strains, comprising acallosal mouse strain I/LnJ, C57BL/6 and BALB/cJ (Schroeter et al., 2017), all strains demonstrated bilateral stimulus-evoked fMRI responses to unilateral hind paw stimulation, thus ruling out minimizing the contribution of transcallosal structural communication as a reason for bilateral FC. Emergence of inter-hemispherical homotopic cortical as well as striato-cortical connectivity was shown to be primarily due to monosynaptic connections (Grandjean et al., 2017); whereas, certain distributed cortical (e.g. default mode network) and subcortical networks emerge through polysynaptic connections in C57BL/6 brains. In BALB/cJ strain, compensation for inter-hemispherical cross-talk could be achieved through the thicker middle portion of cc or possibly through polysynaptic routes. In a rodent model of partial callosotomy (Zhou et al., 2014), initial decrease in inter-hemispherical connectivity was reversed over time, likely due to compensation by remaining axonal pathways. In this study, similar to our findings, an increased intra-hemispherical connectivity was also noted.

#### 4.2.1 Default mode-like network

FC differences were also found in the DMN-like patterns of two mice populations. The DMN maps highlighted larger patterns of connectivity between RSP and parts of TH, CP and dhc areas in the C57BL/6N animals, while the ACA and RSP connect better along the top cortical line and TEa in BALB/cJ. Stronger local connectivity of ACA was noted in the frontal regions of C57Bl/6N brains, in agreement with locally increased density of fibers and higher FA in these areas. Weaker synchrony of the BOLD rsfMRI signal within DMN was previously reported in the C57BL/6 animals (Shah et al., 2016b), when compared to BALB/c and SJL mice. In the present work, we didn’t reproduce the lower C57BL/6N functional connectivity features for DMN when compared with the BALB/cJ strain. We rather observed variance in the spatial organization of this network, with stronger cortico-subcortical communication in C57BL/6N and less functionally connected ACA-RSP. Strain-dependent impact of the medetomidine anesthesia cannot be excluded in this context and depends – among other factors - on the expression levels of alpha-2 adrenoreceptors across the brain.

#### 4.2.2 Functional connectivity signatures of reward aversion pathways

Reward system has consistently been implicated in several pathologies especially ASD and mood disorders (Hägele et al., 2015; Russo and Nestler, 2013) and is relevant for the behavioral phenotypes previously described in the C57BL/6N and BALB/c strains. For instance, BALB/c genetic background seems to favor strong response to stress conditions, show high anxiety and reduced social interaction (Anderzhanova et al., 2013; Brodkin, 2007; Moy et al., 2007; Ohl et al., 2001; Panksepp and Lahvis, 2007). Such phenotypes might stem from vulnerability of specific connectional pathways between key nodes of the reward-aversion system in which ACB is one of the major players. Our results thus demonstrated large patterns of stronger ACB FC in the C57BL/6N mice, including reinforced ACB - VTA dopaminergic pathways and stronger ACB – prefrontal cortical subcortical septal regions.

Activity dynamics of VTA - ACB projection encodes and predicts key features of social interaction in mice. Optogenetic control of cells specifically contributing to this projection is sufficient to modulate social behavior, mediated by dopamine receptor type 1 signaling downstream from VTA to ACB (Gunaydin et al., 2014). Weaker ACB-VTA communication might therefore form the neural substrate of the social reward deficiency previously described in the BALB/c mice. Similarly, differences of directional connectivity of the frontal cingulate (ACA) and striatal limbic network (CP, ACB and AMY) could account for behavioral differences in rewarding effects of addictive substances such as cocaine (Belzung and Barreau, 2000; Crawley et al., 1997). In light of distinct FC patterns we observed in the BALB/cJ mice, it could be argued that specific functioning of reward system determines the atypical behavioral phenotype in this strain.

## 5 Conclusion

Taken together, our findings demonstrate distinct structural and functional brain architectures in two frequently used mouse strains: C57BL/6N and BALB/cJ. In particular, BALB/cJ strain was characterized by marked intra-strain variability; thus, this variability should be taken into account while interpreting results from studies using BALB/cJ. C57BL/6N and BALB/cJ further show divergent brain wide FC diagrams; an essential aspect to be considered in experimental disease models that would also reflect inherent strain differences.

## 7 List of Abbreviations

AAA-CEA-MEA: Anterior/Central /Medial Amygdala
ACA: Anterior Cingulate Area
ACB/NAc: Nucleus accumbens
AD: Axial Diffusivity
AI: Agranular Insula
AIC: Akaike Information Criterion
AMBA: Allen Mouse Brain Atlas
AMY: Amygdala
ANTs: Advanced Normalization Tools
ASD: Autism Spectrum Disorders
AUD: Auditory Area
BOLD: Blood Oxygen-Dependent Signal
BST, BNST: Bed Nucleus of Stria Terminalis
CA: Cornu Ammonis; Ammon’s Horn
cc: Corpus Callosum
CLA: Claustrum
COA-PAA-TR: Cortical/Piriform Amygdala-Postpiriform Transition Area
Cp: Cerebral Peduncle
CP/CPu: Caudate-Putamen
DCA: Directional Connectivity Analysis
DG: Dentate Gyrus
dHIP: Dorsal Hippocampus
DMN: Default Mode Network
DTI: Diffusion Tensor Imaging
ECT: Ectorhinal Area
ENT: Entorhinal Area
EP: Endopiriform Nucleus
fa: Anterior Forceps
FA: Fractional Anisotropy
FC: Functional Connectivity
FD: Fiber Density
FDR: False Discovery Rate
fMRI: Functional Magnetic Resonance Imaging
FOV: Field of View
fr: Fasciculus retroflexus
FMR1: Fragile X Mental Retardation 1 gene
FMR1-KO: Fragile X Mental Retardation 1 gene knockout
FRP: Frontal Pole
FWHM: Full Width at Half Maximum FXS Fragile X syndrome
gcc: Genu of Corpus Callosum
GCM: Group comparison matrix
GE-EPI: Gradient Echo-Echo Planar Imaging
GM: Gray Matter
GU: Gustatory Area
hc: Hippocampus
hrFM: High Resolution Fiber Mapping
HY: Hypothalamus
LA-BLA-BMA-PA: Lateral/Basolateral/Basomedial/Posterior Amygdala
LSX-MSC-TRS: Lateral/Medial Septal Complex-Triangular Nucleus of Septum
LV: Lateral Ventricle
Mbsat: Behavioral State Related Midbrain
mcc: Middle Corpus Callosum
MD^1^: Mean Diffusivity
MD^2^: Medetomidine
mfb: Medial Forebrain Bundle
MH-LH: Medial/Lateral Habenula
MO: Motor Area
MOl: Left Motor Area
Mop: Primary Motor Area
MOr: Right Motor Area
MOs: Secondary Motor Area
mPFC: Medial Prefrontal Cortex
MRI: Magnetic Resonance Imaging
MRN: Midbrain Reticular Nuclei
MVGC: Multivariate Granger Causality Toolbox
NAc/ACB: Nucleus Accumbens
ORB: Orbital Area
PAG: Periaqueductal Gray
PALd: Dorsal Pallidum
PALv: Ventral Pallidum
PAR-POST-PRE-SUB: Para/ Post/Pre-Subiculum-Subiculum
PC: Partial correlation
PERI: Perirhinal Area
PIR: Piriform Area
PL: Prelimbic Area
PRN: Pontine Reticular Nuclei
PTLp: Posterior Parietal Association Area
RARE: Rapid Acquisition with Refocused Echoes
RD: Radial Diffusivity
ROI: Region of interest
rs-fMRI: Resting State Functional Magnetic Resonance Imaging
RSP: Retrosplenial Area
s.c.: Subcutaneous
scc: Splenium of Corpus Callosum
SN: Substantia Nigra
SPM: Statistical Parametric Mapping
spO_2_: Peripheral Oxygen Saturation
SS: Somatosensory Area
SSl: Left Somatosensory Area
SSr: Right Somatosensory Area
SyN: Symmetric Normalization Algorithm
Tcp: Thalamo-cortical Pathways
TE: Echo Time
TeA: Temporal Association Area
TH: Thalamus
TR: Repetition Time
VBM: Voxel-based Morphometry
VBQ: Voxel-based Quantification
vHIP: Ventral Hippocampus
VIS: Visual Area
VISC: Visceral Area
VTA: Ventral Tegmental area
WM: White Matter

---

**Supplementary Figure 1. Group-specific fiber density (FD) maps reveal structural strain differences. (A)** Mean FD for C57BL/6N group. **(B)** Mean FD for BALB/cJ group. **(C-D)** Voxel-wise statistical group comparison of FD maps (p< 0.05, after false discovery rate (FDR) p-value correction; color scales represent t-values) indicate areas of differences **(C)** Contrast C57BL/6N > BALB/cJ: higher fiber density in C57BL/6N mice vs. BALB/cJ group. (D) Contrast BALB/cJ > C57BL/6N: higher fiber density in BALB/cJ mice vs. C57BL/6N group.[Abbreviations: Genu (gcc), middle (mcc) and splenium (scc) of corpus callosum(cc); fasciculus retroflexus(fr), cerebral peduncle(cp), striato-cortical pathways (scp), prelimbic area (PL), anterior cingulate area (ACA), caudate-putamen (CP), temporal association areas (TeA), thalamus (TH), periaqueductal gray (PAG), midbrain reticular nucleus (MRN), motor areas (MO), lateral ventricle(LV).]

## References

Alexander, A.L., Lee, J.E., Lazar, M., Boudos, R., DuBray, M.B., Oakes, T.R., Miller, J.N., Lu, J., Jeong, E.-K., McMahon, W.M., Bigler, E.D., Lainhart, J.E., 2007. Diffusion tensor imaging of the corpus callosum in Autism. NeuroImage 34, 61–73. https://doi.org/10.1016/j.neuroimage.2006.08.032

Anderzhanova, E.A., Bächli, H., Buneeva, O.A., Narkevich, V.B., Medvedev, A.E., Thoeringer, C.K., Wotjak, C.T., Kudrin, V.S., 2013. Strain differences in profiles of dopaminergic neurotransmission in the prefrontal cortex of the BALB/C vs. C57Bl/6 mice: Consequences of stress and afobazole. Eur. J. Pharmacol. 708, 95–104. https://doi.org/10.1016/j.ejphar.2013.03.015

Avants, B.B., Tustison, N.J., Song, G., Cook, P.A., Klein, A., Gee, J.C., 2011. A reproducible evaluation of ANTs similarity metric performance in brain image registration. Neuroimage 54, 2033–2044. https://doi.org/10.1016/j.neuroimage.2010.09.025

Bardella, G., Bifone, A., Gabrielli, A., Gozzi, A., Squartini, T., 2016. Hierarchical organization of functional connectivity in the mouse brain: a complex network approach. Sci. Rep. 6, 32060. https://doi.org/10.1038/srep32060

Barnett, L., Seth, A.K., 2014. The MVGC multivariate Granger causality toolbox: A new approach to Granger-causal inference. J. Neurosci. Methods 223, 50–68. https://doi.org/10.1016/j.jneumeth.2013.10.018

Belzung, C., Barreau, S., 2000. Differences in drug-induced place conditioning between BALB/c and C57B1/6 mice. Pharmacol. Biochem. Behav. 65, 419–423. https://doi.org/10.1016/S0091-3057(99)00212-9

Belzung, C., Griebel, G., 2001. Measuring normal and pathological anxiety-like behaviour in mice: a review. Behav. Brain Res. 125, 141–149.

Biswal, B., Zerrin Yetkin, F., Haughton, V.M., Hyde, J.S., 1995. Functional connectivity in the motor cortex of resting human brain using echo-planar mri. Magn. Reson. Med. 34, 537–541.

Bonelli, R.M., Cummings, J.L., 2007. Frontal-subcortical circuitry and behavior. Dialogues Clin. Neurosci. 9, 141–151.

Bray, S., Hirt, M., Jo, B., Hall, S.S., Lightbody, A.A., Walter, E., Chen, K., Patnaik, S., Reiss, A.L., 2011. Aberrant Frontal Lobe Maturation in Adolescents with Fragile X Syndrome is Related to Delayed Cognitive Maturation. Biol. Psychiatry, Genetic and Environmental Contributors to Disturbed Cortical Development in Developmental Disorders 70, 852–858. https://doi.org/10.1016/j.biopsych.2011.05.038

Brodkin, E.S., 2007. BALB/c mice: Low sociability and other phenotypes that may be relevant to autism. Behav. Brain Res., Animal Models for Autism 176, 53–65. https://doi.org/10.1016/j.bbr.2006.06.025

Brodkin, E.S., Hagemann, A., Nemetski, S.M., Silver, L.M., 2004. Social approach–avoidance behavior of inbred mouse strains towards DBA/2 mice. Brain Res. 1002, 151–157. https://doi.org/10.1016/j.brainres.2003.12.013

Calamante, F., Tournier, J.-D., Heidemann, R.M., Anwander, A., Jackson, G.D., Connelly, A., 2011. Track density imaging (TDI): Validation of super resolution property. NeuroImage 56, 1259–1266. https://doi.org/10.1016/j.neuroimage.2011.02.059

Calcagno, E., Canetta, A., Guzzetti, S., Cervo, L., Invernizzi, R.W., 2007. Strain differences in basal and post-citalopram extracellular 5-HT in the mouse medial prefrontal cortex and dorsal hippocampus: relation with tryptophan hydroxylase-2 activity. J. Neurochem. 103, 1111–1120. https://doi.org/10.1111/j.1471-4159.2007.04806.x

Chen, C., Kim, J.J., Thompson, R.F., Tonegawa, S., 1996. Hippocampal lesions impair contextual fear conditioning in two strains of mice. Behav. Neurosci. 110, 1177–1180. https://doi.org/10.1037/0735-7044.110.5.1177

Chuang, K.-H., Nasrallah, F.A., 2017. Functional networks and network perturbations in rodents. NeuroImage 163, 419–436. https://doi.org/10.1016/j.neuroimage.2017.09.038

Clipperton-Allen, A.E., Ingrao, J.C., Ruggiero, L., Batista, L., Ovari, J., Hammermueller, J., Armstrong, J.N., Bienzle, D., Choleris, E., Turner, P.V., 2015. Long-Term Provision of Environmental Resources Alters Behavior but not Physiology or Neuroanatomy of Male and Female BALB/c and C57BL/6 Mice. J. Am. Assoc. Lab. Anim. Sci. 54, 718–730.

Crawley, J.N., Belknap, J.K., Collins, A., Crabbe, J.C., Frankel, W., Henderson, N., Hitzemann, R.J., Maxson, S.C., Miner, L.L., Silva, A.J., 1997. Behavioral phenotypes of inbred mouse strains: implications and recommendations for molecular studies. Psychopharmacology (Berl.) 132, 107–124.

de Sá-Calçada, D., Roque, S., Branco, C., Monteiro, S., Cerqueira-Rodrigues, B., Sousa, N., Palha, J.A., Correia-Neves, M., 2015. Exploring Female Mice Interstrain Differences Relevant for Models of Depression. Front. Behav. Neurosci. 9. https://doi.org/10.3389/fnbeh.2015.00335

Dennis, E.L., Thompson, P.M., 2013a. Mapping connectivity in the developing brain. Int. J. Dev. Neurosci. 31, 525–542. https://doi.org/10.1016/j.ijdevneu.2013.05.007

Dennis, E.L., Thompson, P.M., 2013b. Typical and atypical brain development: a review of neuroimaging studies. Dialogues Clin. Neurosci. 15, 359–384.

Dodero, L., Damiano, M., Galbusera, A., Bifone, A., Tsaftsaris, S.A., Scattoni, M.L., Gozzi, A., 2013. Neuroimaging Evidence of Major Morpho-Anatomical and Functional Abnormalities in the BTBR T+TF/J Mouse Model of Autism. PLOS ONE 8, e76655. https://doi.org/10.1371/journal.pone.0076655

Draganski, B., Ashburner, J., Hutton, C., Kherif, F., Frackowiak, R.S.J., Helms, G., Weiskopf, N., 2011. Regional specificity of MRI contrast parameter changes in normal ageing revealed by voxel-based quantification (VBQ). NeuroImage 55, 1423–1434. https://doi.org/10.1016/j.neuroimage.2011.01.052

Eagle, D.M., Baunez, C., Hutcheson, D.M., Lehmann, O., Shah, A.P., Robbins, T.W., 2008. Stop-Signal Reaction-Time Task Performance: Role of Prefrontal Cortex and Subthalamic Nucleus. Cereb. Cortex 18, 178–188. https://doi.org/10.1093/cercor/bhm044

Ellegood, J., Anagnostou, E., Babineau, B.A., Crawley, J.N., Lin, L., Genestine, M., DiCicco-Bloom, E., Lai, J.K.Y., Foster, J.A., Peñagarikano, O., Geschwind, D.H., Pacey, L.K., Hampson, D.R., Laliberté, C.L., Mills, A.A., Tam, E., Osborne, L.R., Kouser, M., Espinosa-Becerra, F., Xuan, Z., Powell, C.M., Raznahan, A., Robins, D.M., Nakai, N., Nakatani, J., Takumi, T., van Eede, M.C., Kerr, T.M., Muller, C., Blakely, R.D., Veenstra-VanderWeele, J., Henkelman, R.M., Lerch, J.P., 2015. Clustering autism: using neuroanatomical differences in 26 mouse models to gain insight into the heterogeneity. Mol. Psychiatry 20, 118–125. https://doi.org/10.1038/mp.2014.98

Ellegood, J., Babineau, B.A., Henkelman, R.M., Lerch, J.P., Crawley, J.N., 2013. Neuroanatomical analysis of the BTBR mouse model of autism using magnetic resonance imaging and diffusion tensor imaging. NeuroImage 70, 288–300. https://doi.org/10.1016/j.neuroimage.2012.12.029

Ellegood, J., Pacey, L.K., Hampson, D.R., Lerch, J.P., Henkelman, R.M., 2010. Anatomical phenotyping in a mouse model of fragile X syndrome with magnetic resonance imaging. NeuroImage, Imaging Genetics 53, 1023–1029. https://doi.org/10.1016/j.neuroimage.2010.03.038

Fairless, A.H., Dow, H.C., Kreibich, A.S., Torre, M., Kuruvilla, M., Gordon, E., Morton, E.A., Tan, J., Berrettini, W.H., Li, H., Abel, T., Brodkin, E.S., 2012. Sociability and brain development in BALB/cJ and C57BL/6J mice. Behav. Brain Res. 228, 299–310. https://doi.org/10.1016/j.bbr.2011.12.001

Fairless, A.H., Dow, H.C., Toledo, M.M., Malkus, K.A., Edelmann, M., Li, H., Talbot, K., Arnold, S.E., Abel, T., Brodkin, E.S., 2008. Low sociability is associated with reduced size of the corpus callosum in the BALB/cJ inbred mouse strain. Brain Res. 1230, 211–217. https://doi.org/10.1016/j.brainres.2008.07.025

Fairless, A.H., Katz, J.M., Vijayvargiya, N., Dow, H.C., Kreibich, A.S., Berrettini, W.H., Abel, T., Brodkin, E.S., 2013. Development of home cage social behaviors in BALB/cJ vs. C57BL/6J mice. Behav. Brain Res. 237, 338–347. https://doi.org/10.1016/j.bbr.2012.08.051

Fontaine, D.A., Davis, D.B., 2016. Attention to Background Strain Is Essential for Metabolic Research: C57BL/6 and the International Knockout Mouse Consortium. Diabetes 65, 25–33. https://doi.org/10.2337/db15-0982

Fox, A.S., Shelton, S.E., Oakes, T.R., Converse, A.K., Davidson, R.J., Kalin, N.H., 2010. Orbitofrontal Cortex Lesions Alter Anxiety-Related Activity in the Primate Bed Nucleus of Stria Terminalis. J. Neurosci. 30, 7023–7027. https://doi.org/10.1523/JNEUROSCI.5952-09.2010

Frazier, T.W., Youngstrom, E.A., Speer, L., Embacher, R., Law, P., Constantino, J., Findling, R.L., Hardan, A.Y., Eng, C., 2012. Validation of Proposed DSM-5 Criteria for Autism Spectrum Disorder. J. Am. Acad. Child Adolesc. Psychiatry 51, 28–40.e3. https://doi.org/10.1016/j.jaac.2011.09.021

Good, C.D., Johnsrude, I.S., Ashburner, J., Henson, R.N.A., Friston, K.J., Frackowiak, R.S.J., 2001. A Voxel-Based Morphometric Study of Ageing in 465 Normal Adult Human Brains. NeuroImage 14, 21–36. https://doi.org/10.1006/nimg.2001.0786

Grandjean, J., Zerbi, V., Balsters, J., Wenderoth, N., Rudina, M., 2017. The structural basis of large-scale functional connectivity in the mouse. J. Neurosci. 0438–17. https://doi.org/10.1523/JNEUROSCI.0438-17.2017

Grange, P., 2018. Topology of the mesoscale connectome of the mouse brain. ArXiv181104698 Q-Bio.

Grubb, S.C., Churchill, G.A., Bogue, M.A., 2004. A collaborative database of inbred mouse strain characteristics. Bioinformatics 20, 2857–2859. https://doi.org/10.1093/bioinformatics/bth299

Gunaydin, L.A., Grosenick, L., Finkelstein, J.C., Kauvar, I.V., Fenno, L.E., Adhikari, A., Lammel, S., Mirzabekov, J.J., Airan, R.D., Zalocusky, K.A., Tye, K.M., Anikeeva, P., Malenka, R.C., Deisseroth, K., 2014. Natural Neural Projection Dynamics Underlying Social Behavior. Cell 157, 1535–1551. https://doi.org/10.1016/j.cell.2014.05.017

Guzzetti, S., Calcagno, E., Canetta, A., Sacchetti, G., Fracasso, C., Caccia, S., Cervo, L., Invernizzi, R.W., 2008. Strain differences in paroxetine-induced reduction of immobility time in the forced swimming test in mice: role of serotonin. Eur. J. Pharmacol. 594, 117–124. https://doi.org/10.1016/j.ejphar.2008.07.031

Hägele, C., Schlagenhauf, F., Rapp, M., Sterzer, P., Beck, A., Bermpohl, F., Stoy, M., Ströhle, A., Wittchen, H.-U., Dolan, R.J., Heinz, A., 2015. Dimensional psychiatry: reward dysfunction and depressive mood across psychiatric disorders. Psychopharmacology (Berl.) 232, 331–341. https://doi.org/10.1007/s00213-014-3662-7

Harsan, L.-A., Dávid, C., Reisert, M., Schnell, S., Hennig, J., von Elverfeldt, D., Staiger, J.F., 2013. Mapping remodeling of thalamocortical projections in the living reeler mouse brain by diffusion tractography. Proc. Natl. Acad. Sci. U. S. A. 110, E1797–1806. https://doi.org/10.1073/pnas.1218330110

Horsfield, M.A., Jones, D.K., 2002. Applications of diffusion-weighted and diffusion tensor MRI to white matter diseases – a review. NMR Biomed. 15, 570–577. https://doi.org/10.1002/nbm.787

Hübner, N.S., Mechling, A.E., Lee, H.-L., Reisert, M., Bienert, T., Hennig, J., von Elverfeldt, D., Harsan, L.-A., 2017. The connectomics of brain demyelination: Functional and structural patterns in the cuprizone mouse model. NeuroImage 146, 1–18. https://doi.org/10.1016/j.neuroimage.2016.11.008

Ingalhalikar, M., Smith, A., Parker, D., Satterthwaite, T.D., Elliott, M.A., Ruparel, K., Hakonarson, H., Gur, R.E., Gur, R.C., Verma, R., 2014. Sex differences in the structural connectome of the human brain. Proc. Natl. Acad. Sci. 111, 823–828. https://doi.org/10.1073/pnas.1316909110

Jacome, L.F., Burket, J.A., Herndon, A.L., Deutsch, S.I., 2011. Genetically inbred Balb/c mice differ from outbred Swiss Webster mice on discrete measures of sociability: relevance to a genetic mouse model of autism spectrum disorders. Autism Res. 4, 393–400. https://doi.org/10.1002/aur.218

Jones, D.K., 2004. The effect of gradient sampling schemes on measures derived from diffusion tensor MRI: A Monte Carlo study. Magn. Reson. Med. 51, 807–815. https://doi.org/10.1002/mrm.20033

Just, M.A., Cherkassky, V.L., Keller, T.A., Kana, R.K., Minshew, N.J., 2006. Functional and Anatomical Cortical Underconnectivity in Autism: Evidence from an fMRI Study of an Executive Function Task and Corpus Callosum Morphometry. Cereb. Cortex 17, 951–961. https://doi.org/10.1093/cercor/bhl006

Keane, T.M., Goodstadt, L., Danecek, P., White, M.A., Wong, K., Yalcin, B., Heger, A., Agam, A., Slater, G., Goodson, M., Furlotte, N.A., Eskin, E., Nellåker, C., Whitley, H., Cleak, J., Janowitz, D., Hernandez-Pliego, P., Edwards, A., Belgard, T.G., Oliver, P.L., McIntyre, R.E., Bhomra, A., Nicod, J., Gan, X., Yuan, W., Weyden, L. van der, Steward, C.A., Bala, S., Stalker, J., Mott, R., Durbin, R., Jackson, I.J., Czechanski, A., Guerra-Assunção, J.A., Donahue, L.R., Reinholdt, L.G., Payseur, B.A., Ponting, C.P., Birney, E., Flint, J., Adams, D.J., 2011. Mouse genomic variation and its effect on phenotypes and gene regulation. Nature 477, 289. https://doi.org/10.1038/nature10413

Kim, S., Pickup, S., Fairless, A.H., Ittyerah, R., Dow, H.C., Abel, T., Brodkin, E.S., Poptani, H., 2012. Association between sociability and diffusion tensor imaging in BALB/cJ mice: DIFFUSION TENSOR IMAGING AND SOCIABILITY. NMR Biomed. 25, 104–112. https://doi.org/10.1002/nbm.1722

Knox, J.E., Harris, K.D., Graddis, N., Whitesell, J.D., Zeng, H., Harris, J.A., Shea-Brown, E., Mihalas, S., 2018. High resolution data-driven model of the mouse connectome. bioRxiv 293019. https://doi.org/10.1101/293019

Kumar, M., Kim, S., Pickup, S., Chen, R., Fairless, A.H., Ittyerah, R., Abel, T., Brodkin, E.S., Poptani, H., 2012. Longitudinal in-vivo diffusion tensor imaging for assessing brain developmental changes in BALB/cJ mice, a model of reduced sociability relevant to autism. Brain Res. 1455, 56–67. https://doi.org/10.1016/j.brainres.2012.03.041

Lai, J.K.Y., Lerch, J.P., Doering, L.C., Foster, J.A., Ellegood, J., 2016. Regional brain volumes changes in adult male FMR1-KO mouse on the FVB strain. Neuroscience 318, 12–21. https://doi.org/10.1016/j.neuroscience.2016.01.021

Lassi, G., Tucci, V., 2017. Gene-environment interaction influences attachment-like style in mice. Genes Brain Behav. 16, 612–618. https://doi.org/10.1111/gbb.12385

Le Bihan, D., 2014. Diffusion MRI: what water tells us about the brain. EMBO Mol. Med. 6, 569–573. https://doi.org/10.1002/emmm.201404055

Le Bihan, D., Breton, E., 1985. Imagerie de diffusion in-vivo par résonance magnétique nucléaire. Comptes-Rendus Académie Sci. 93, 27–34.

Lein, E.S., Hawrylycz, M.J., Ao, N., Ayres, M., Bensinger, A., Bernard, A., Boe, A.F., Boguski, M.S., Brockway, K.S., Byrnes, E.J., Chen, Lin, Chen, Li, Chen, T.-M., Chi Chin, M., Chong, J., Crook, B.E., Czaplinska, A., Dang, C.N., Datta, S., Dee, N.R., Desaki, A.L., Desta, T., Diep, E., Dolbeare, T.A., Donelan, M.J., Dong, H.-W., Dougherty, J.G., Duncan, B.J., Ebbert, A.J., Eichele, G., Estin, L.K., Faber, C., Facer, B.A., Fields, R., Fischer, S.R., Fliss, T.P., Frensley, C., Gates, S.N., Glattfelder, K.J., Halverson, K.R., Hart, M.R., Hohmann, J.G., Howell, M.P., Jeung, D.P., Johnson, R.A., Karr, P.T., Kawal, R., Kidney, J.M., Knapik, R.H., Kuan, C.L., Lake, J.H., Laramee, A.R., Larsen, K.D., Lau, C., Lemon, T.A., Liang, A.J., Liu, Y., Luong, L.T., Michaels, J., Morgan, J.J., Morgan, R.J., Mortrud, M.T., Mosqueda, N.F., Ng, L.L., Ng, R., Orta, G.J., Overly, C.C., Pak, T.H., Parry, S.E., Pathak, S.D., Pearson, O.C., Puchalski, R.B., Riley, Z.L., Rockett, H.R., Rowland, S.A., Royall, J.J., Ruiz, M.J., Sarno, N.R., Schaffnit, K., Shapovalova, N.V., Sivisay, T., Slaughterbeck, C.R., Smith, S.C., Smith, K.A., Smith, B.I., Sodt, A.J., Stewart, N.N., Stumpf, K.-R., Sunkin, S.M., Sutram, M., Tam, A., Teemer, C.D., Thaller, C., Thompson, C.L., Varnam, L.R., Visel, A., Whitlock, R.M., Wohnoutka, P.E., Wolkey, C.K., Wong, V.Y., Wood, M., Yaylaoglu, M.B., Young, R.C., Youngstrom, B.L., Feng Yuan, X., Zhang, B., Zwingman, T.A., Jones, A.R., 2007. Genome-wide atlas of gene expression in the adult mouse brain. Nature 445, 168–176. https://doi.org/10.1038/nature05453

Liska, A., Galbusera, A., Schwarz, A.J., Gozzi, A., 2015. Functional connectivity hubs of the mouse brain. NeuroImage 115, 281–291. https://doi.org/10.1016/j.neuroimage.2015.04.033

Liska, A., Gozzi, A., 2016. Can Mouse Imaging Studies Bring Order to Autism Connectivity Chaos? Front. Neurosci. 10. https://doi.org/10.3389/fnins.2016.00484

McFarlane, H.G., Kusek, G.K., Yang, M., Phoenix, J.L., Bolivar, V.J., Crawley, J.N., 2008. Autism-like behavioral phenotypes in BTBR T+tf/J mice. Genes Brain Behav. 7, 152–163. https://doi.org/10.1111/j.1601-183X.2007.00330.x

Mechling, A.E., Arefin, T., Lee, H.-L., Bienert, T., Reisert, M., Ben Hamida, S., Darcq, E., Ehrlich, A., Gaveriaux-Ruff, C., Parent, M.J., Rosa-Neto, P., Hennig, J., von Elverfeldt, D., Kieffer, B.L., Harsan, L.-A., 2016. Deletion of the mu opioid receptor gene in mice reshapes the reward–aversion connectome. Proc. Natl. Acad. Sci. 201601640. https://doi.org/10.1073/pnas.1601640113

Miller, V.M., Gupta, D., Neu, N., Cotroneo, A., Boulay, C.B., Seegal, R.F., 2013. Novel inter-hemispheric white matter connectivity in the BTBR mouse model of autism. Brain Res. 1513, 26–33. https://doi.org/10.1016/j.brainres.2013.04.001

Moy, S.S., Nadler, J.J., Young, N.B., Perez, A., Holloway, L.P., Barbaro, R.P., Barbaro, J.R., Wilson, L.M., Threadgill, D.W., Lauder, J.M., Magnuson, T.R., Crawley, J.N., 2007. Mouse behavioral tasks relevant to autism: Phenotypes of 10 inbred strains. Behav. Brain Res, Animal Models for Autism 176, 4–20. https://doi.org/10.1016/j.bbr.2006.07.030

Oh, S.W., Harris, J.A., Ng, L., Winslow, B., Cain, N., Mihalas, S., Wang, Q., Lau, C., Kuan, L., Henry, A.M., Mortrud, M.T., Ouellette, B., Nguyen, T.N., Sorensen, S.A., Slaughterbeck, C.R., Wakeman, W., Li, Y., Feng, D., Ho, A., Nicholas, E., Hirokawa, K.E., Bohn, P., Joines, K.M., Peng, H., Hawrylycz, M.J., Phillips, J.W., Hohmann, J.G., Wohnoutka, P., Gerfen, C.R., Koch, C., Bernard, A., Dang, C., Jones, A.R., Zeng, H., 2014. A mesoscale connectome of the mouse brain. Nature 508, 207–214. https://doi.org/10.1038/nature13186

Ohl, F., Sillaber, I., Binder, E., Keck, M.E., Holsboer, F., 2001. Differential analysis of behavior and diazepam-induced alterations in C57BL/6N and BALB/c mice using the modified hole board test. J. Psychiatr. Res. 35, 147–154. https://doi.org/10.1016/S0022-3956(01)00017-6

Palumbo, M.L., Canzobre, M.C., Pascuan, C.G., Ríos, H., Wald, M., Genaro, A.M., 2010. Stress induced cognitive deficit is differentially modulated in BALB/c and C57Bl/6 mice: Correlation with Th1/Th2 balance after stress exposure. J. Neuroimmunol. 218, 12–20. https://doi.org/10.1016/j.jneuroim.2009.11.005

Panksepp, J.B., Lahvis, G.P., 2007. Social reward among juvenile mice. Genes Brain Behav. 6, 661–671.

Peng, D.X., Kelley, R.G., Quintin, E.-M., Raman, M., Thompson, P.M., Reiss, A.L., 2014. Cognitive and behavioral correlates of caudate subregion shape variation in fragile X syndrome. Hum. Brain Mapp. 35, 2861–2868. https://doi.org/10.1002/hbm.22376

Pervolaraki, E., Tyson, A.L., Pibiri, F., Poulter, S.L., Reichelt, A.C., Rodgers, R.J., Clapcote, S.J., Lever, C., Andreae, L.C., Dachtler, J., 2019. The within-subject application of diffusion tensor MRI and CLARITY reveals brain structural changes in Nrxn2 deletion mice. Mol. Autism 10, 8. https://doi.org/10.1186/s13229-019-0261-9

Pilz, L.K., Quiles, C.L., Dallegrave, E., Levandovski, R., Hidalgo, M.P.L., Elisabetsky, E., Pilz, L.K., Quiles, C.L., Dallegrave, E., Levandovski, R., Hidalgo, M.P.L., Elisabetsky, E., 2015. Differential susceptibility of BALB/c, C57BL/6N, and CF1 mice to photoperiod changes. Rev. Bras. Psiquiatr. 37, 185–190. https://doi.org/10.1590/1516-4446-2014-1454

Qiu, M., Ye, Z., Li, Q., Liu, G., Xie, B., Wang, J., 2011. Changes of Brain Structure and Function in ADHD Children. Brain Topogr. 24, 243–252. https://doi.org/10.1007/s10548-010-0168-4

Reisert, M., Mader, I., Anastasopoulos, C., Weigel, M., Schnell, S., Kiselev, V., 2011. Global fiber reconstruction becomes practical. NeuroImage 54, 955–962. https://doi.org/10.1016/j.neuroimage.2010.09.016

Roebroeck, A., Formisano, E., Goebel, R., 2005. Mapping directed influence over the brain using Granger causality and fMRI. NeuroImage 25, 230–242. https://doi.org/10.1016/j.neuroimage.2004.11.017

Russo, S.J., Nestler, E.J., 2013. The brain reward circuitry in mood disorders. Nat. Rev. Neurosci. 14, 609–625. https://doi.org/10.1038/nrn3381

Sankoorikal, G.M.V., Kaercher, K.A., Boon, C.J., Lee, J.K., Brodkin, E.S., 2006. A Mouse Model System for Genetic Analysis of Sociability: C57BL/6J Versus BALB/cJ Inbred Mouse Strains. Biol. Psychiatry 59, 415–423. https://doi.org/10.1016/j.biopsych.2005.07.026

Schroeter, A., Grandjean, J., Schlegel, F., Saab, B.J., Rudin, M., 2017. Contributions of structural connectivity and cerebrovascular parameters to functional magnetic resonance imaging signals in mice at rest and during sensory paw stimulation. J. Cereb. Blood Flow Metab. 37, 2368–2382. https://doi.org/10.1177/0271678X16666292

Shah, D., Blockx, I., Keliris, G.A., Kara, F., Jonckers, E., Verhoye, M., Van der Linden, A., 2016a. Cholinergic and serotonergic modulations differentially affect large-scale functional networks in the mouse brain. Brain Struct. Funct. 221, 3067–3079. https://doi.org/10.1007/s00429-015-1087-7

Shah, D., Deleye, S., Verhoye, M., Staelens, S., Van der Linden, A., 2016b. Resting-state functional MRI and [18F]-FDG PET demonstrate differences in neuronal activity between commonly used mouse strains. NeuroImage 125, 571–577. https://doi.org/10.1016/j.neuroimage.2015.10.073

Shepherd, G.M.G., 2013. Corticostriatal connectivity and its role in disease. Nat. Rev. Neurosci. 14, 278–291. https://doi.org/10.1038/nrn3469

Stouffer, S.A., Suchman, E.A., DeVinney, L.C., Star, S.A., Williams, R.M.J., 1949. Adjustment During Army Life. Princet. NJ Princet. Univ. Press.

Tustison, N.J., Avants, B.B., Cook, P.A., Yuanjie Zheng Egan, A., Yushkevich, P.A., Gee, J.C., 2010. N4ITK: Improved N3 Bias Correction. IEEE Trans. Med. Imaging 29, 1310–1320. https://doi.org/10.1109/TMI.2010.2046908

Wahlsten, D., 1974. Heritable aspects of anomalous myelinated fibre tracts in the forebrain of the laboratory mouse. Brain Res. 68, 1–18. https://doi.org/10.1016/0006-8993(74)90530-7

Watson, G.D.R., Smith, J.B., Alloway, K.D., 2017. Interhemispheric connections between the infralimbic and entorhinal cortices: The endopiriform nucleus has limbic connections that parallel the sensory and motor connections of the claustrum. J. Comp. Neurol. 525, 1363–1380. https://doi.org/10.1002/cne.23981

Weible, A.P., 2013. Remembering to attend: The anterior cingulate cortex and remote memory. Behav. Brain Res. 245, 63–75. https://doi.org/10.1016/j.bbr.2013.02.010

Wu, D., Zhang, J., 2016. In vivo mapping of macroscopic neuronal projections in the mouse hippocampus using high-resolution diffusion MRI. NeuroImage 125, 84–93. https://doi.org/10.1016/j.neuroimage.2015.10.051

Wu, G.-R., Liao, W., Stramaglia, S., Ding, J.-R., Chen, H., Marinazzo, D., 2013. A blind deconvolution approach to recover effective connectivity brain networks from resting state fMRI data. Med. Image Anal. 17, 365–374. https://doi.org/10.1016/j.media.2013.01.003

Yochum, C.L., Medvecky, C.M., Cheh, M.A., Bhattacharya, P., Wagner, G.C., 2010. Differential development of central dopaminergic and serotonergic systems in BALB/c and C57BL/6J mice. Brain Res. 1349, 97–104. https://doi.org/10.1016/j.brainres.2010.06.031

Yoshida, K., Mimura, Y., Ishihara, R., Nishida, H., Komaki, Y., Minakuchi, T., Tsurugizawa, T., Mimura, M., Okano, H., Tanaka, K.F., Takata, N., 2016. Physiological effects of a habituation procedure for functional MRI in awake mice using a cryogenic radiofrequency probe. J. Neurosci. Methods 274, 38–48. https://doi.org/10.1016/j.jneumeth.2016.09.013

Zhang, J., Aggarwal, M., Mori, S., 2012. Structural insights into the rodent CNS via diffusion tensor imaging. Trends Neurosci. 35, 412–421. https://doi.org/10.1016/j.tins.2012.04.010

Zhang, X., Li, Q., Wong, N., Zhang, M., Wang, W., Bu, B., McAlonan, G.M., 2015. Behaviour and prefrontal protein differences in C57BL/6N and 129 X1/SvJ mice. Brain Res. Bull. 116, 16–24. https://doi.org/10.1016/j.brainresbull.2015.05.003

Zhou, I.Y., Liang, Y.-X., Chan, R.W., Gao, P.P., Cheng, J.S., Hu, Y., So, K.-F., Wu, E.X., 2014. Brain resting-state functional MRI connectivity: Morphological foundation and plasticity. NeuroImage 84, 1–10. https://doi.org/10.1016/j.neuroimage.2013.08.037

