## Supplementary figures and images for "Mapping the living mouse brain neural architecture: strain specific patterns of brain structural and functional connectivity"

### Supplementary Figure 1

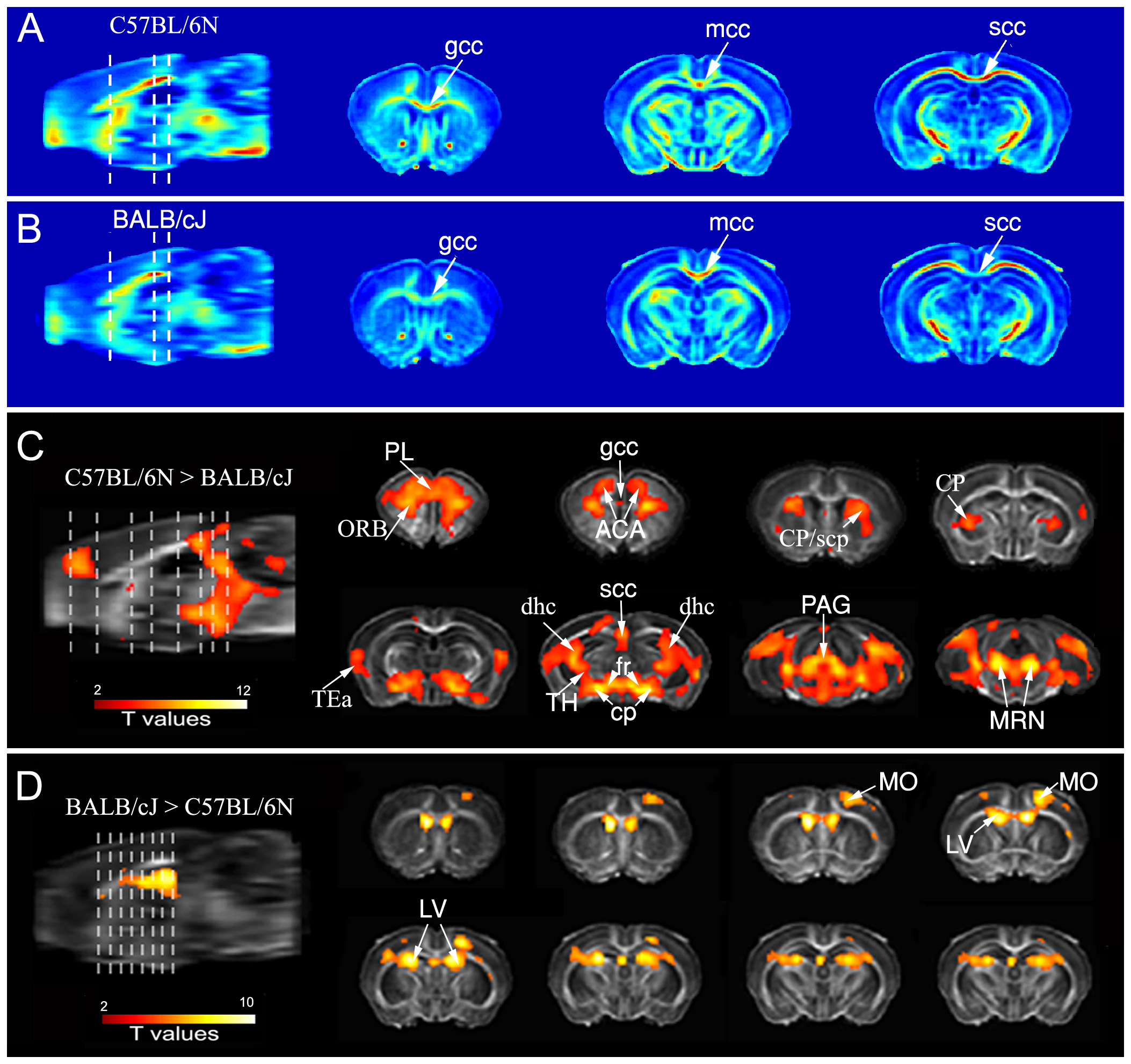
